# The EspN transcription factor is an infection-dependent regulator of the ESX-1 system in *M. marinum*

**DOI:** 10.1101/2023.02.15.528779

**Authors:** Kathleen R. Nicholson, Rachel M. Cronin, Aruna R. Menon, Madeleine K. Jennisch, David M. Tobin, Patricia A. Champion

## Abstract

Bacterial pathogens use protein secretion systems to translocate virulence factors into the host and to control bacterial gene expression. The ESX-1 (ESAT-6 system 1) secretion system facilitates disruption of the macrophage phagosome during infection, enabling access to the cytoplasm, and regulates widespread gene expression in the mycobacterial cell. The transcription factors contributing to the ESX-1 transcriptional network during mycobacterial infection are not known. We showed that the EspM and WhiB6 transcription factors regulate the ESX-1 transcriptional network *in vitro* but are dispensable for macrophage infection by *Mycobacterium marinum*. In this study, we used our understanding of the ESX-1 system to identify EspN, a critical transcription factor that controls expression of the ESX-1 genes during infection, but whose effect is not detectable under standard laboratory growth conditions. Under laboratory conditions, EspN activity is masked by the EspM repressor. In the absence of EspM, we found that EspN is required for ESX-1 function because it activates expression of the *whiB6* transcription factor gene, and specific ESX-1 substrate and secretory component genes. Unlike the other transcription factors that regulate ESX-1, EspN is required for *M. marinum* growth within and cytolysis of macrophages, and for disease burden in a zebrafish larval model of infection. These findings demonstrate that EspN is an infection-dependent regulator of the ESX-1 transcriptional network, which is essential for mycobacterial pathogenesis. Moreover, our findings suggest that ESX-1 expression is controlled by a genetic switch that responds to host specific signals.

**Importance:** Pathogenic mycobacteria cause acute and long-term diseases, including human tuberculosis. The ESX-1 system transports proteins that control the host response to infection and promotes bacterial survival. Although ESX-1 transports proteins, it also controls gene expression in the bacteria. In this study, we identify an undescribed transcription factor that controls the expression of ESX-1 genes, and is required for both macrophage and animal infection. However, this transcription factor is not the primary regulator of ESX-1 genes under standard laboratory conditions. These findings identify a critical transcription factor that controls expression of a major virulence pathway during infection, but whose effect is not detectable with standard laboratory strains and growth conditions.

## Introduction

In diverse pathogens, secretion systems transport virulence factor substrates that promote bacterial survival in different niches within the host (1, 2). Pathogenic mycobacteria use conserved ESAT-6 secretion (ESX) systems to facilitate disruption of the macrophage phagosome during infection, enabling access to the cytoplasm and toxification of the macrophage (3–5). *Mycobacterium tuberculosis*, the causative agent of human tuberculosis, and *Mycobacterium marinum*, a poikilothermic fish pathogen, share a conserved ESX-1 secretion system that is required for pathogenesis (6–8). The ESX-1 system is composed of ESX core components (Ecc’s) that reside within the mycobacterial cell membrane (CM) (7, 9, 10). During macrophage infection, the ESX-1 system transports protein substrates that are secreted components of the ESX-1 machinery (11) and effectors that promote phagosomal damage (4, 5, 12, 13). Phagosomal damage enables the essential interaction between pathogenic mycobacteria and the macrophage cytoplasm (5, 14–17).

Protein secretion systems in several pathogens also regulate bacterial gene expression (2, 18, 19). In addition to transporting proteins, we and others have shown that ESX-1 controls gene expression globally in *M. marinum* and *M. tuberculosis* (20–22). Loss of the ESX-1 membrane complex leads to widespread changes in gene expression (20, 21). The gene most responsive to loss of the ESX-1 membrane complex, *whiB6*, encodes a transcription factor that activates ESX-1 substrate gene expression (20, 23, 24). We showed that EspM is a direct repressor of *whiB6* gene transcription in the absence of the ESX-1 membrane complex (20, 22). Reduced WhiB6 leads to lower levels of ESX-1 substrate gene transcription, preventing accumulation of ESX-1 substrates in the absence of the transport machinery (20). Genetic deletion of the *espM* gene, which is divergently encoded from the *whiB6* gene, leads to derepression of *whiB6* expression, and increased levels of ESX-1 substrates in *M. marinum* during *in vitro* growth (22).

Despite regulating ESX-1 substrate gene expression under laboratory conditions, neither WhiB6 nor EspM is essential for virulence in macrophage models of infection (20, 22). Therefore, ESX-1 may be differentially regulated in the laboratory and at different points or niches during infection. We previously used *lacZ+* transcriptional fusions to measure the transcriptional activity of the *M. marinum whiB6* promoter in the presence and absence of the EspM and WhiB6 transcription factors. In the absence of both EspM and WhiB6, there was a significant increase in *β*-galactosidase activity, ~2-fold higher than in the absence of EspM alone (22). These data suggested that in the absence of both WhiB6 and EspM, *whiB6* gene expression was activated. We exploited our knowledge of a negative regulator of ESX-1 expression, EspM, as a tool to dissect the genetic circuitry that regulates the ESX-1 regulatory network (Fig. 1A).

**Figure 1.**
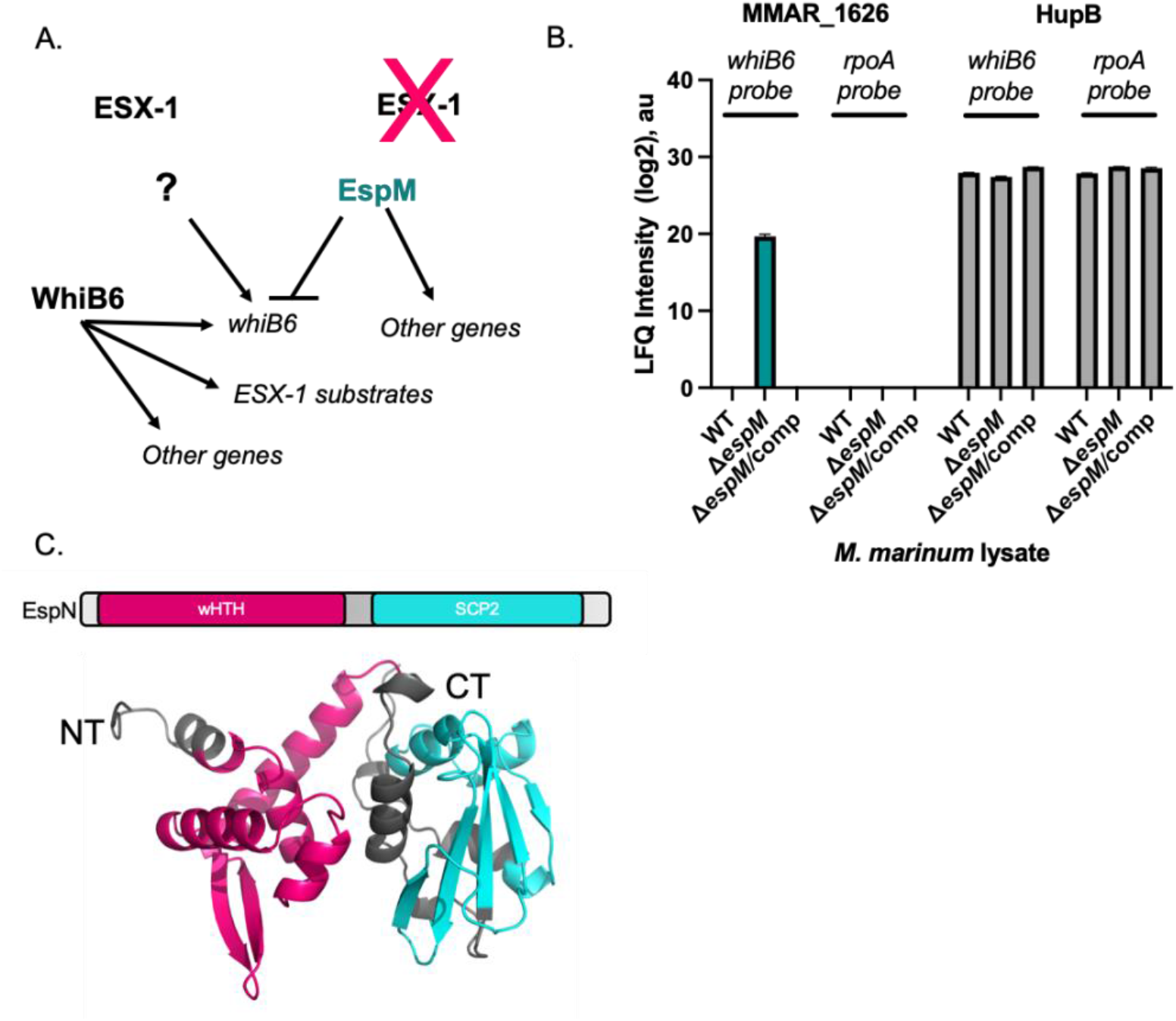
EspN is a putative transcription factor that binds the *whiB6* promoter. **A.** Schematic of the ESX-1 transcriptional network. In the presence of the ESX-1 system, WhiB6 is positively autoregulated, and positively regulates the expression of the ESX-1 substrates, and other genes. In the absence of the ESX-1 system, EspM directly represses *whiB6* gene expression, reducing ESX-1 substrate levels. In the absence of WhiB6 and EspM, an unknown activator upregulates *whiB6* expression (?). **B.** Mass spectrometry analysis of the DNA affinity chromatography showing enrichment of the MMAR_1626 and HupB proteins. The HupB protein binds nonspecifically to both DNA probes. The scale represents the log2 intensity of MS peak arears. a.u. arbitrary units. The data was published in Sanchez et al (22). **C.** The predicted domain structure of MMAR_1626, which we renamed EspN. Modeled using RoseTTAFold from Robetta (80). Model confidence: 0.80. wHTH: winged helix turn helix, SCP2: sterol carrier protein 2.

## Results

### EspN is a putative transcription factor that binds the *whiB6* promoter

To identify regulators responsible for activation of the *whiB6* promoter in the absence of EspM we used the *whiB6-espM* intergenic region to enrich proteins from lysate generated from the Δ*espM M. marinum* strain. Mass spectrometry identified MMAR_1626 as the only protein enriched for binding the *whiB6-espM* intergenic region in the absence of *espM* (Fig. 1B) (22).

MMAR_1626 is an unstudied putative transcriptional regulator with a predicted N-terminal winged helix-turn-helix domain, and a C-terminal sterol carrier protein 2 (SCP2) domain (Fig. 1C) (25, 26). *MMAR_1626* gene orthologs are found across *Mycobacterium* and other high G+C Gram-positive bacteria, including members of the TB complex, *M. tuberculosis (Rv1725c*) and *M. bovis (Mb1754c*), and non-pathogens, including *M. smegmatis* (27, 28). Based on the following data, we propose naming this protein EspN, according to accepted nomenclature (29).

### EspN is an activator of *whiB6* gene expression in the absence of EspM

Our data led us to hypothesize that EspN is an activator of *whiB6* gene expression (Fig. 1A). To test this hypothesis, we generated an unmarked *espN* deletion strain using allelic exchange (Fig. S1A and S1B) (30). The parental *M. marinum* M strain (wild type, WT) has a *whiB6* allele with a C-terminal 3xFLAG epitope integrated at the *whiB6* locus (20). We generated the corresponding complementation strain by integrating a copy of the *espN* gene behind the constitutive mycobacterial optimal promoter at the *attL* site. We tested *espN* expression in these strains using qRT-PCR (Fig. S2A). We detected the *espN* transcript in the WT strain, but not in the Δ*espN* strain (*P*=.0002, compared to the WT strain, inset). *espN* was significantly overexpressed (> 40-fold) in the Δ*espN/pespN* strain as compared to the WT strain (*P*<.0114, Fig S2A).

To evaluate the role of EspN on *whiB6* expression, we measured *whiB6* transcript levels in the Δ*espN* and Δ*espN/pespN* strains relative to the WT strain and Δ*eccCb_1_* strains using qRT-PCR (Fig. 2A). The Δ*eccCb_1_* strain does not produce the EccCb_1_ protein, a cytoplasmic component of the ESX-1 system that directly interacts with the membrane components (7). The loss of EccCb_1_ results in a loss of the entire ESX-1 membrane complex (10, 20, 31), and results in repression of *whiB6* gene expression (20, 22). *whiB6* expression in the Δ*eccCb_1_* strain was significantly reduced compared to the WT strain (*P*<.0001) due to a loss of the ESX-1 membrane complex (20). In contrast, *whiB6* expression in the Δ*espN* and Δ*espN/pespN* strains was stochastic and not significantly different from the WT strain. These findings were confirmed using western blot analysis on *M. marinum* cell-associated proteins (Fig. 2B). We detected WhiB6-Fl protein in the WT strain (Fig. 2B, lane 1). Consistent with our previous data, deletion of the *eccCb_1_* gene resulted in a loss of detectable WhiB6-Fl protein (Fig. 2B, lane 2). The levels of WhiB6-Fl protein in the Δ*espN* strain or the Δ*espN*/p*espN* complementation strains were similar to the WT strain (Fig. 2B, lanes 3 and 4). From these data, we conclude that there was no EspN-dependent change in *whiB6* expression in *M. marinum* under the conditions tested.

**Figure 2.**
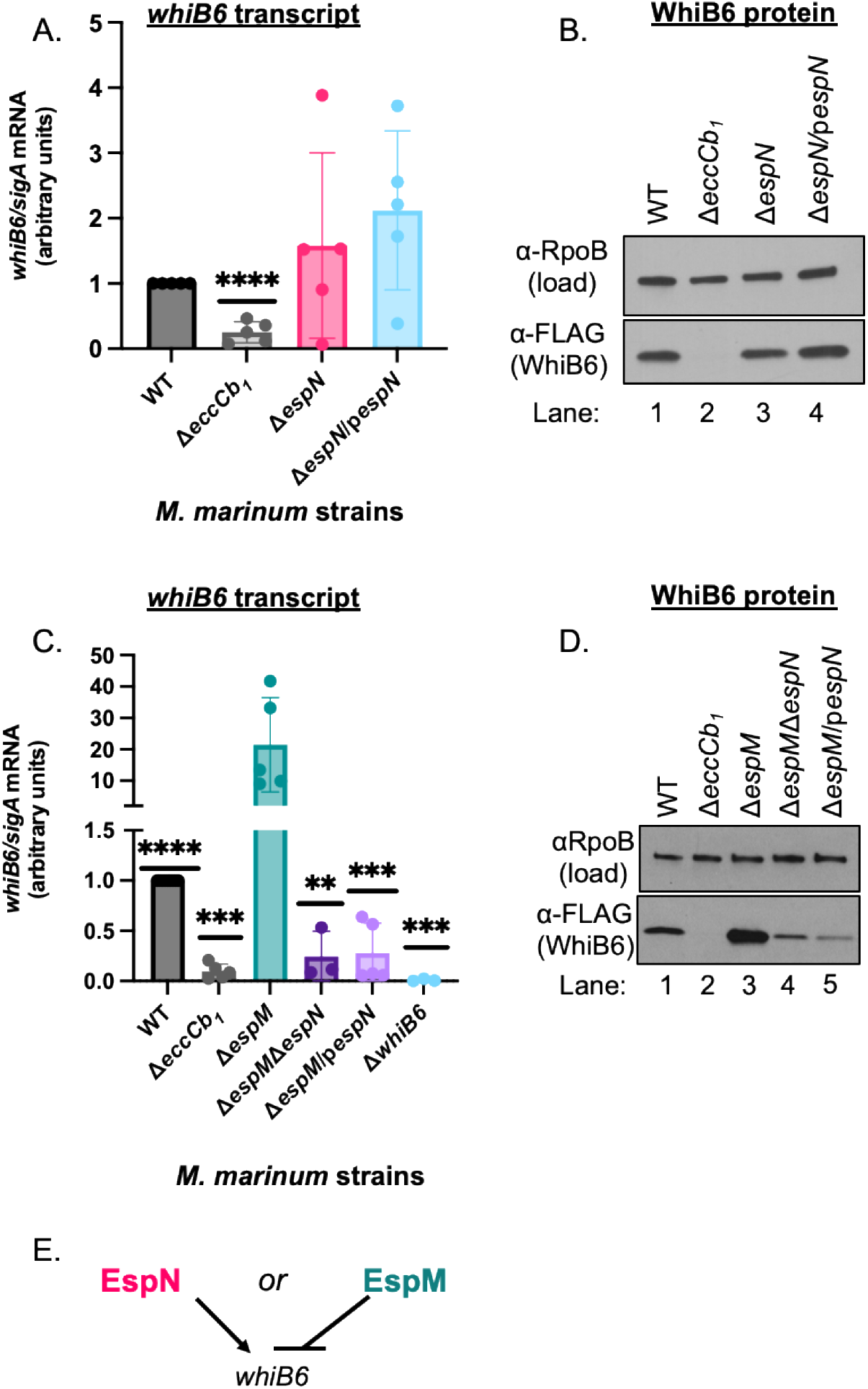
EspN activates *whiB6* expression in the absence of EspM. **A.** Relative qRT-PCR analysis of *M. marinum* strains compared to *sigA* transcript levels. Statistical analysis was performed using a one-way ANOVA (* *P*=.0359) followed by a Dunnett’s multiple comparisons test relative to the WT strain. Significance between the WT and Δ*eccCb_1_* strain was determined using an unpaired students t-test. **** *P*<.0001. **B.** Western blot analysis of *M. marinum* cell-associated proteins. RpoB is a loading control. All strains include a *whiB6-3xFl* allele at the *whiB6* locus (20). **C.** qRT-PCR of the *whiB6* transcript relative to *sigA*. Significance was determined using a one-way ordinary ANOVA (*P*=.0001), followed by a Tukey’s multiple comparison test. Significance shown is relative to the Δ*EspM* strain, with additional statistics of interest discussed in the text. **** *P*<.0001, *** *P*=.0002 for Δ*EccCb_1_, ** P*=.0010, *** *P*=.0002 for Δ*espM/pespN*, *** *P*=.0009 for Δ*whiB6*. **D.** Western blot analysis of 10μg of *M. marinum* whole cell lysates. RpoB serves as a loading control. All strains include a *whiB6-3xFl* allele at the *whiB6* locus (20) **E**. Schematic summarizing findings from Figure 2.

We identified both EspM, the repressor of *whiB6* expression, and EspN using DNA affinity chromatography with the same *whiB6* promoter DNA (22). While we identified EspM specifically bound to the *whiB6* promoter following incubation with WT *M. marinum* lysate, the EspN protein was only specifically bound to the *whiB6* promoter following incubation with Δ*espM* lysate (22). We hypothesized that EspN may regulate *whiB6* expression in the absence of and in opposition to the EspM repressor. To test this hypothesis, we deleted the *espN* gene in a Δ*espM M. marinum* strain. We also introduced an integrating plasmid constitutively expressing the *espN* gene into the Δ*espM* strain. We confirmed the loss and restoration of *espN* expression using qRT-PCR (Fig. S2B). The *espN* transcript was absent from the Δ*espMΔespN* strain, and significantly increased in the Δ*espM/pespN* strain (~60-80-fold, *P*<.0001). We performed qRT-PCR to quantify changes in *whiB6* transcript levels in the *espN* deletion and overexpression strains compared to the Δ*espM* strain (Fig. 2C). *whiB6* was expressed in the WT strain and was significantly reduced in the Δ*eccCb_1_* strain relative to the WT strain (*P*<.0001) (20, 22). *whiB6* expression was significantly increased in the Δ*espM* strain (*P*<.0001), consistent with the loss of the EspM repressor (22). Both genetic deletion and overexpression of the *espN* gene in the Δ*espM* strain resulted in a significant reduction in *whiB6* transcript levels compared to Δ*espM* strain (*P*=.0002 and *P*=.0009). The levels of *whiB6* in both strains were also significantly reduced compared to the WT strain (*P*<.0001). We interpret these data to mean that overexpression of *espN* in the Δ*espM* strain results in a loss of EspN function, similar to the deletion strain (32). Moreover, we conclude that EspN is required for the elevated levels of WhiB6 in the Δ*espM* strain.

To confirm these findings, we performed a western blot analysis on cell-associated fractions of *M. marinum* mutants to detect changes in WhiB6 protein (Fig. 2D). Similar to Figure 2B and our previous work, the WhiB6-Fl protein was produced to detectable levels in the WT strain and reduced in the Δ*eccCb_1_* strain (lanes 1 and 2, Fig. 2D). WhiB6-Fl protein levels were increased in the Δ*espM* strain (lane 3), consistent with the role of EspM as a *whiB6* repressor. Deletion or overexpression of *espN* in the absence of EspM resulted in a loss of WhiB6-Fl protein (lanes 4 and 5, Fig. 2D), consistent with the loss of *whiB6* expression in Fig. 2C. Together, these data suggest that EspN activates the expression of *whiB6* in the absence of EspM (Fig. 2E), working in opposition to EspM at the *whiB6* promoter. Moreover, overexpression of the *espN* gene results in a loss of function of EspN in the absence of EspM.

### EspN is required for ESX-1 dependent lytic activity

*M. marinum* exhibit contact-dependent hemolytic activity that requires the ESX-1 system. To test if EspN was required for ESX-1 function in the absence of EspM, we measured the membranolytic activity of the relevant *M. marinum* strains by measuring sheep red blood cell (sRBC) lysis (Fig. 3A). Water and PBS were “no bacteria” controls. They resulted in total sRBC lysis and baseline sRBC lysis as measured by OD405, respectively (Fig. 3A). *M. marinum* lysed sRBCs in an ESX-1-dependent manner (Δ*eccCb_1_* vs WT, *P*<.0001). Deletion or overexpression of the *espN* gene did not significantly impact the hemolytic activity of *M. marinum* relative to the WT strain. From these data we conclude that EspN is dispensable for ESX-1-mediated hemolysis in the presence of EspM. The Δ*espM* strain lysed sRBCs to a slightly lower level than the WT strain (*P*=.0122). The deletion or overexpression of *espN* in the Δ*espM* strain abrogated hemolysis (*P*<.0001), similar to the Δ*eccCb_1_* strain. From these data, we conclude that EspN is essential for the lytic activity of ESX-1 in the absence of EspM.

**Figure 3.**
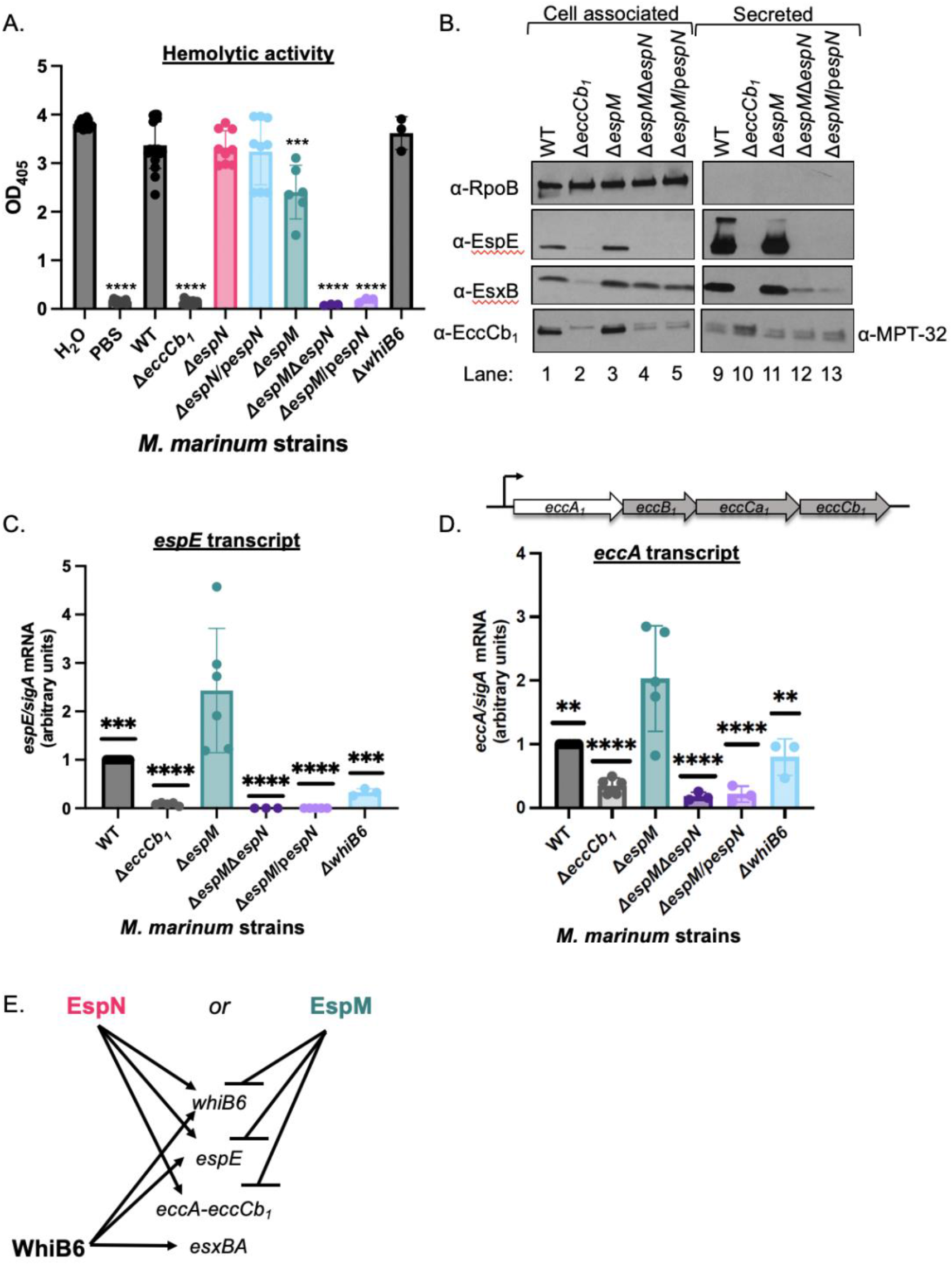
EspN activates expression of ESX-1 components and substrates. **A.** Sheep red blood cell lysis (sRBC) measuring hemolytic activity of *M. marinum*. Statistical analysis was performed using a one-way ordinary ANOVA followed by a Dunnett’s multiple comparisons test relative to the WT strain. **** *P*<0.0001, *** *P*=.0003. **B.** Western blot of 10μg of *M. marinum* cell-associated proteins. All strains include a *whiB6-Fl* allele. RpoB is a control for lysis. MPT-32 is a loading control for the secreted fractions. **C.** Relative qRT analysis the *espE* transcript compared to *sigA* transcript levels in *M. marinum*. Statistical analysis was performed using a one-way ordinary ANOVA (*P*<.0001) followed by a Tukey’s multiple comparison test. Significance is shown relative to the Δ*espM* strain. *** *P*=0.0010 (WT), **** *P*<.0001, *** *P*=0.0003 (Δ*whiB6*). **D.** Relative qRT analysis of the *eccA* transcript compared to *sigA* transcript levels in *M. marinum*. Statistical analysis was performed using a one-way ordinary ANOVA (*P*<.0001) followed by a Tukey’s multiple comparison test. Significance shown relative to the Δ*espM* strain. ** *P*=.0012 (WT), *P*=.0026 (Δ*whiB6*), **** *P*<.0001, *** *P*=0.0003. **E.** Schematic summarizing the results of Figure 3, and the contributions of EspN, EspM and WhiB6 to regulating ESX-1 gene expression.

### EspN is a transcriptional activator of genes encoding for ESX-1 components and substrates

We sought to determine why EspN was necessary for ESX-1 hemolytic activity. WhiB6 regulates the expression of ESX-1 substrates (20, 23, 24). If EspN was essential for lytic activity solely because EspN activated *whiB6* expression, then we would expect that the hemolytic activity of the Δ*espMΔespN* strain would phenocopy the Δ*whiB6* strain. Consistent with our previous findings, the Δ*whiB6* strain retained hemolytic activity (20). These data indicate that the loss of WhiB6 alone in the Δ*espMΔespN* strain is insufficient to explain the loss of hemolytic activity.

We reasoned that the loss of hemolytic activity could be due to loss of ESX-1 secretion (8). To test this idea, we performed western blot analysis to measure the production and secretion of two ESX-1 substrates, EsxB and EspE. EsxB is secreted with EsxA as an early substrate of the ESX-1 system, and is likely a secreted component of the ESX-1 system in the mycolate outer membrane (7, 11). The loss of EsxB results in a loss of hemolytic activity because it abrogates secretion of the ESX-1 substrates (8, 33). EspE is a late substrate of the ESX-1 system (11). The loss of EspE results in a loss of hemolytic activity, but does not impact the secretion of other ESX-1 substrates, except EspF (11, 33). As shown in Figure 3B, EsxB and EspE were produced in the WT strain (lane 1, αEsxB, αEspE) and secreted into the culture supernatant during *in vitro* growth (lane 9). EsxB and EspE proteins were reduced in the Δ*eccCb_1_* strain (lane 2), due to repression of *whiB6* expression by EspM (22). Neither protein was secreted into the culture supernatant (lane 10, Fig. 3B) due to a loss of the ESX-1 membrane complex. Both EspE and EsxB were produced (lane 3, Fig. 3B) and secreted from the Δ*espM* strain (lane 11, Fig. 3B), consistent with our prior findings (22). Deletion or overexpression of the *espN* gene in the Δ*espM* strain resulted in a loss of EspE production (lanes 4 and 5, Fig. 3B). Although EsxB was produced in both strains, the secretion of EsxB was reduced compared to the WT and Δ*espM* strains (lanes 12 and 13, Fig. 3B).

Reduced EsxB secretion could be due to reduced levels of the ESX-1 membrane complex. EccCb_1_ is a cytoplasmic component of the ESX-1 system that directly interacts with the membrane components (7). The loss of EccCb_1_ results in a loss of the entire ESX-1 membrane complex (20). We measured EccCb_1_ levels by western blot analysis (Fig. 3B). EccCb_1_ protein was detected in the cell-associated proteins from the WT strain and absent from the Δ*eccCb_1_* strain (lanes 1 and 2, lower band, Fig. 3B). The Δ*espM* strain produced EccCb_1_ protein (lane 3, Fig. 3B). In the absence of EspM and EspN, EccCb_1_ protein levels were reduced to levels lower than the WT strain (lanes 4 and 5, Fig. 3B).

Based on the findings in Figure 3B, we tested if EspN regulated the expression of *espE* or the membrane complex in the absence of EspM. To determine if loss of EspE protein was due to decreased *espE* expression, we performed qRT-PCR (Fig. 3C). Deletion of the *eccCb_1_* gene led to a significant reduction in *espE* expression relative to the WT strain (*P*<.0001), consistent with Fig. 4A and our previous findings (22, 34). Deletion of *espM* significantly increased *espE* gene expression relative to the WT strain (*P*=.0010)(22). These data confirm that EspM represses the expression of the *espE* gene in *M. marinum*, consistent with our prior RNA sequencing results (22). Deletion or overexpression of *espN* in the Δ*espM* strain resulted a complete loss of *espE* gene expression, significantly lower than both the Δ*espM* (*P*<.0001) and the WT strains (*P*<.0001). From these data we conclude that EspN is required for expression of *espE* in the Δ*espM* strain (Fig. 3E).

**Figure 4.**
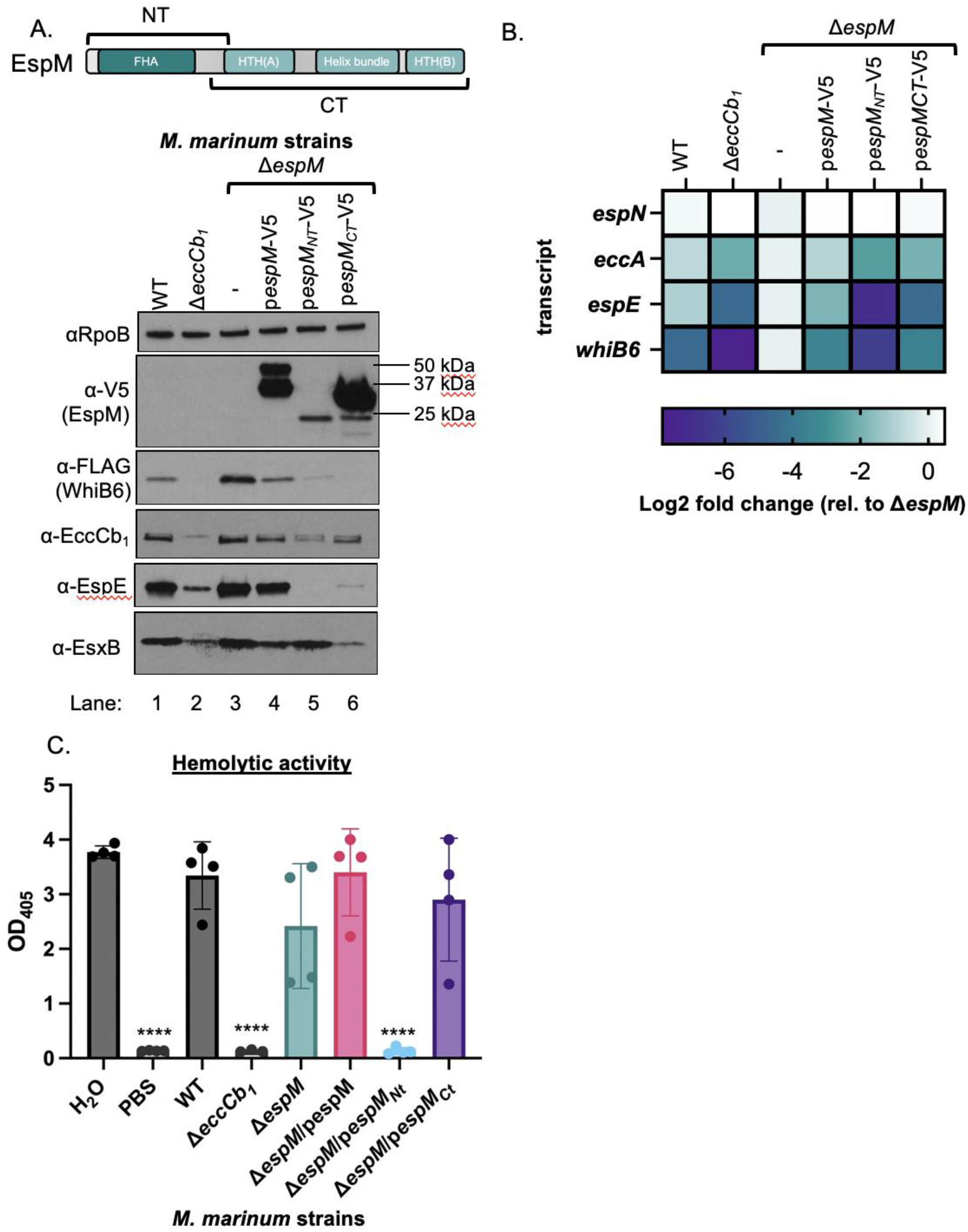
The EspM N-terminus negatively regulates EspN. **A.** Western blot of *M. marinum* whole cell lysates. All strains include a *whiB6-Fl* allele. EspM complementation strains are tagged at the C-terminus with a V5 tag. RpoB is a loading control. EspE is an ESX-1 substrate. 10μg of protein was loaded in each lane. **B.** Relative qRT-PCR analysis of *M. marinum* strains compared to *sigA* transcript levels. The values were normalized to those in the Δ*espM* strain and plotted as the log2 fold change relative to the Δ*espM* strain as a heat map. The qRT-PCR data plotted in this heat map are shown in Fig. S5A-D. **C.** Sheep red blood cell lysis (sRBC) measuring hemolytic activity of *M. marinum*. Statistical analysis was performed using a one-way ANOVA (*P*<.0001) followed by a Dunnett’s multiple comparisons test (**** *P*<.0001).

WhiB6 positively regulates *espE* gene expression. Importantly, deletion of the *whiB6* gene resulted in a significant reduction (*P*=.0003), but not a loss of the *espE* transcript. Therefore, the loss of WhiB6 from the Δ*espMΔespN* strain was insufficient to cause the measured loss of *espE* expression.

The *eccCb_1_* gene is downstream of genes that encode for three other ESX-1 components (Fig. 3D, *eccA, eccB, eccCa_1_*). It is not known if the *ecc* genes in the ESX-1 locus are operonic. To test if the observed loss of EccCb_1_ protein was due to reduced expression of the *ecc* genes, we measured *eccA* expression using qRT-PCR (Fig. 3D). We measured a significant reduction in the levels of *eccA* in the Δ*eccCb_1_* strain as compared to the WT strain (*P*<.0001). Deletion of *espM* led to a significant increase in *eccA* levels compared to the WT strain (*P*=.0012). Together, the reduction of *eccA* levels in the Δ*eccCb_1_* and the increased levels of *eccA* in the Δ*espM* strain suggest that EspM represses expression of the *eccA* gene (Fig. 3E). The deletion or overexpression of the *espN* gene in the Δ*espM* strain significantly reduced *eccA* expression as compared to the Δ*espM* (*P*<.0001) and the WT strains (*P*<.0001). Interestingly, the levels of *eccA* transcript in the Δ*whiB6* strain, although significantly reduced from the Δ*espM* strain (*P=.0026*), were not significantly different from the WT strain. From these data, we conclude that EspN activates *eccA* gene expression and EspM represses *eccA* gene expression. Moreover, the regulation of *eccA* by EspN is independent of WhiB6 (Fig. 3E).

### The EspM N-terminus post-transcriptionally regulates EspN

Our findings suggest that EspN directly activates *whiB6* expression, and is either a direct or indirect activator of *espE* and *eccA* expression. Importantly, our data show that EspN activates ESX-1-gene expression only in the absence of EspM (Fig. 3E). Therefore, we sought to understand how EspN was connected to our established ESX-1 transcriptional network (Fig. 1A). We first tested if EspM and EspN regulate each other at the transcriptional level. We measured the expression of the *espN* gene in the Δ*espM* strain, and the *espM* gene in the Δ*espN* strain (Fig. S4A and B). We did not observe any significant change in *espN* expression in the Δ*espM* strain, or in *espM* expression in the Δ*espN* strain. From these data, we conclude that EspM and EspN do not regulate each other transcriptionally under the conditions tested.

The EspM protein consists of 4 putative domains, including an N-terminal fork-head associated domain (FHA) and two helix-turn-helix (HTH) domains separated by a helical bundle domain (Fig. 4A) (35). The FHA domain alone (EspMNT in Fig. 4A) does not bind DNA *in vitro* (22). The two HTH domains with the helical bundle (EspMCT in Fig. 4A) are sufficient to bind the *whiB6* promoter *in vitro* and to repress *whiB6* expression in *M. marinum* (22, 35). To further understand the functions of the EspM domains, we expressed the *espM, espMNT* and *espMCT* genes including a C-terminal V5 epitope tag in the Δ*espM* strain. All EspM-V5 proteins were expressed in *M. marinum* (lanes 4-6, Fig. 4A). Both the EspM full-length and EspMCT proteins resulted in multiple bands as detected by the αV5 antibody, consistent with N-terminal processing events. The EspMNT protein resulted in a single size appropriate band (lane 5, Fig. 4A). Consistent with our prior findings, expression of the EspM-V5 protein reduced the WhiB6-Fl protein and significantly reduced the *whiB6* transcript relative to the Δ*espM* strain (lane 4 vs 5, Fig. 4B, and Fig. S5A) (35). Expression of the EspM-V5 protein also resulted in reduced EccCb_1_, EspE, and EsxB protein levels (lane 4) compared to the Δ*espM* strain. We measured significant reductions in the expression of the *eccA* and *espE* genes (Fig. 4B and Fig. S5B and C). Expression of the *espMCT* gene resulted in a loss of detectable WhiB6-Fl protein and further reductions of the EccCb_1_, EspE and EsxB proteins compared to the Δ*espM* and Δ*espM/pespMV5* strains (lanes 6 vs 3 and 4, Fig. 4A). Likewise, we measured significant reductions in the levels of *whiB6, eccA* and *espE* gene expression relative to the Δ*espM* strain. Interestingly, expression of the *espMNT* gene in the Δ*espM* strain resulted in a loss of EspE protein and a reduction of the WhiB6-Fl and EccCb_1_ protein as compared to both the Δ*espM* and the WT strains. Although the EspMNT does not bind the *whiB6* promoter *in vitro* (22), we measured significant reductions in the expression of the *whiB6, espE*, and *eccA* genes compared to the Δ*espM* and complemented strains (Fig. 4B and Fig. S5). None of the strains tested exhibited significant reductions in *espN* expression.

These data suggested that the Δ*espM/pespMNT*-V5 strain effectively phenocopied the Δ*espMΔespN* strain (Fig. 3B). To test if the Δ*espM/espMNT-V5* strain further phenocopied the Δ*espMΔespN* strain, we performed sRBC lysis assays on the strains in Figures 4A and 4B. We reasoned that if the Δ*espM/pespMNT*-V5 phenocopied the Δ*espM*Δ*espN* strain, it would be non-hemolytic (Fig. 3A). Expression of the *espM-V5* or *espMCT-V5* genes in the Δ*espM* strain did not result in a significant change in hemolytic activity (Fig. 4B). However, expression of the *espMNT-V5* abrogated hemolytic activity of the Δ*espM* strain (*P*<.0001). From these data, we conclude that the expression of the EspMNT protein phenocopies the loss of the EspN protein in the Δ*espM* strain. Together our data support a model by which the EspM FHA domain negatively regulates EspN post-transcriptionally.

### EspN is required during *M. marinum* infection of macrophages and zebrafish

Our data support a model in which EspM and EspN function in opposition to regulate WhiB6 and the ESX-1 system (Fig. 3E). EspM represses *whiB6* expression in the absence of the ESX-1 system, and the genes encoding ESX-1 components (22). In the absence of EspM, our data support that EspN activates *whiB6* and ESX-1 gene expression. However, under *in vitro* growth conditions, the presence of EspM obscures the regulatory role of EspN (Fig. 2). We previously found that EspM was dispensable for *M. marinum* virulence in a macrophage model (22). Therefore, we hypothesized that there may be signals governing the switch between EspM and EspN in the host that may be absent during *in vitro* growth.

*M. marinum* cause macrophage cytolysis in an ESX-1-dependent manner. Strains lacking the ESX-1 system are retained in the phagosome and are non-cytolytic (4). We infected with *M. marinum* (MOI of 4) and imaged 24 hours post-infection (hpi) to measure macrophage cytotoxicity using ethidium homodimer (EthD-1) staining (Fig. 5A). EthD-1 is a membrane-impermeable fluorescent molecule that specifically stains DNA in cells with permeabilized cell membranes, reflecting the cytolytic activity of *M. marinum* (8, 36). As shown in Figure 5A, infection with WT *M. marinum* resulted in macrophage cytotoxicity. Uninfected macrophages and macrophages infected with the Δ*eccCb_1_* strain exhibited significantly less cytotoxicity than infection with the WT strain (*P*<.00001). The Δ*espN* strain showed a statistically significant decrease in macrophage cytotoxicity compared to the WT strain, similar to the Δ*eccCb_1_* ESX-1-compromised mutant strain (*P*<.0001). Cytotoxicity was partially complemented by overexpressing *espN* in the Δ*espN* strain (*P*<.0001). The Δ*espMΔespN* and Δ*espM/pespN* strains were non-cytolytic, similar to the Δ*eccCb_1_* strain (Fig. S6).

**Figure 5.**
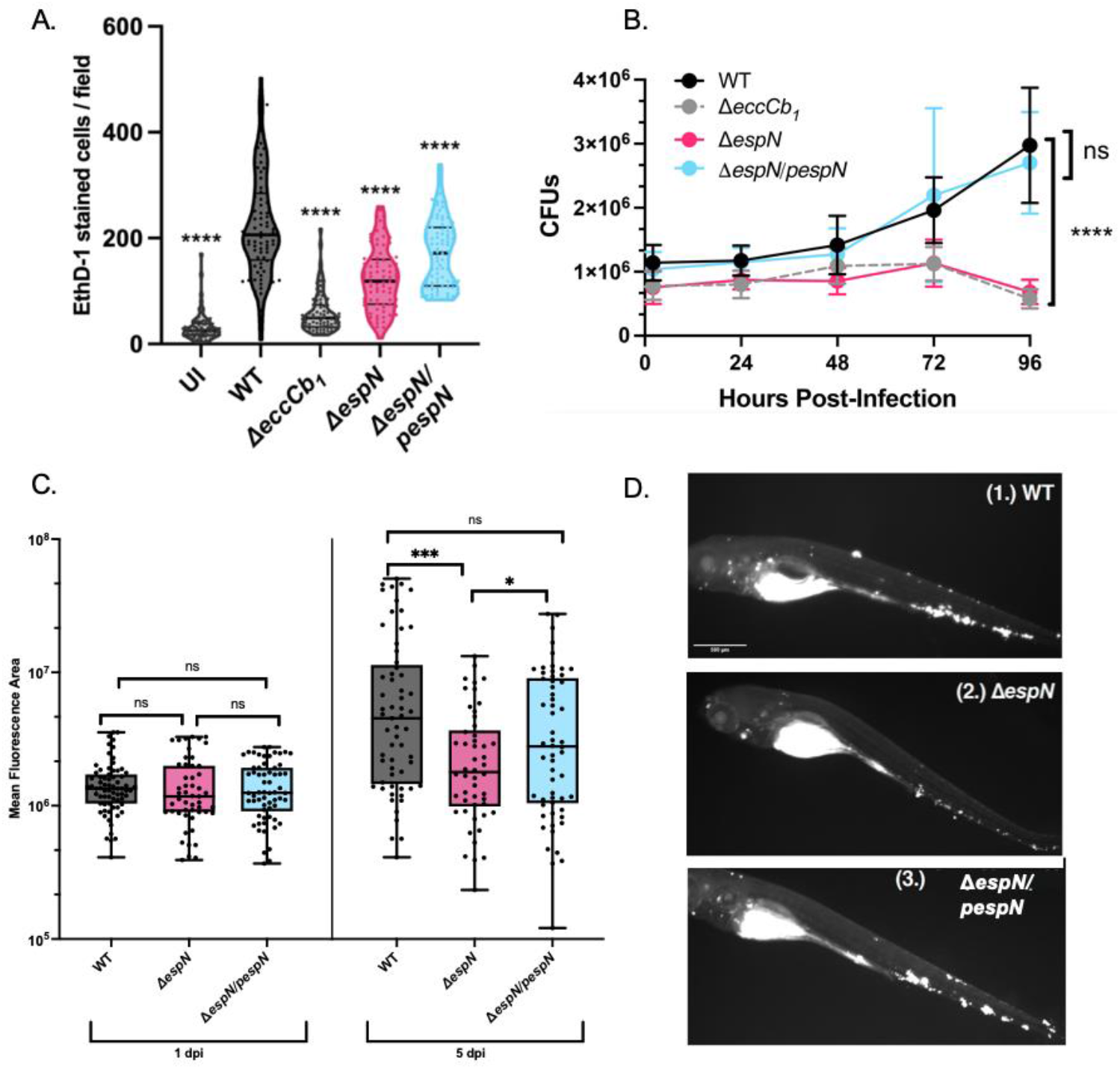
EspN is required for pathogenesis. **A.** Macrophage cytolysis as measured by EthD-1 staining 24h post-infection with *M. marinum* at an MOI of 4. Statistical analysis was performed using a one-way ANOVA followed by a Dunnett’s multiple comparisons test relative to the WT strain. (**** *P*<.0001) Each dot represents the number of EthD-1-stained cells in a single field. A total of 10 fields were counted using ImageJ for each well. Processing of 3 wells was performed for each biological replicate. A total of 90 fields were counted for each strain. **B.** Colony forming units (CFU) of *M. marinum* strains MOI = 0.2. The data points are the average of three independent biological replicates. Significance was determined using an ordinary 2 way ANOVA (*P*<.0001) followed by a Tukey’s multiple comparison test compared to the WT strain. The significance shown is compared to the WT strain at 96-hours post infection (*P*<.0001). The Δ*eccCb_1_* and Δ*espN* strains were also significantly different from the WT strain at 72 hpi (*P*=.0021 and *P*=.0024, respectively). **C.** *M. marinum* burden in zebrafish infection measured using bacterial mCerulean fluorescence. Data are comprised of two biological replicates with 20-30 independent infections per replicate. Statistical analyses were performed using one-way ANOVA followed by a Tukey multiple comparison of each group to the WT strain (*** *P*=.00012; * *P*=.021). **D.** Representative images of zebrafish infected with an initial dose of 150-200 fluorescent bacilli for (1.) WT, (2.) Δ*espN*, or (3.) Δ*espN/pespN* at 5 days post-infection. Scale bar is 500μm.

To test if *espN* was required for mycobacterial growth in macrophages, we performed CFU analysis (Fig. 5B). As published previously, the WT *M. marinum* strain grew within macrophages during the course of the infection. In contrast, the Δ*eccCb_1_* strain was attenuated for growth in macrophages, and resulted in CFUs that were significantly different from the WT strain at 72 and 96 hpi. Consistent with the cytotoxicity data, the Δ*espN* strain was attenuated for growth in the macrophage as compared to the WT strain, similar to the Δ*eccCb_1_* strain. Strikingly, expression of the *espN* gene restored growth of *M. marinum* similar to the WT strain. From these data we conclude that the ESX-1 transcriptional network and EspN are required for *M. marinum* growth and cytotoxicity during macrophage infection.

We next tested if EspN was important for animal infection. To examine the *in vivo* consequences of EspN deletion, we made constitutively fluorescent versions of the WT, Δ*espN*, and complemented *M. marinum* strains. Using the zebrafish larval model of mycobacterial infection, we monitored infection burden over five days of infection as measured by a validated Fluorescence Pixel Count (FPC) assay (Fig. 5C and 5D) (37, 38). Bacterial burden was substantially reduced at five days post-infection. We found that *in vivo, ΔespN* had 3.4-fold reduced burden compared to WT at five days post-infection (*P*=.00012) and was complemented by constitutive expression of the *espN* gene. (Fig. 5C and 5D). From these data, we conclude that EspN is required during zebrafish infection by *M. marinum*.

## Discussion

Our prior studies hinted at the existence of an additional activator of the ESX-1 system (22). We used our knowledge of a negative regulator of ESX-1 expression, EspM, as a tool to dissect the genetic circuitry that regulates the ESX-1 regulatory network *in vivo*, arriving at a novel ESX-1 regulator that specifically functions in the host. We discovered and characterized EspN, an infection-dependent transcriptional regulator of the ESX-1 system. Our data supports that EspN and EspM function as a switch that regulates expression of both the ESX-1 secretion machinery and specific substrates. In the absence of the ESX-1 system, EspM represses the expression of the *whiB6* gene as well as genes encoding the ESX-1 components (EccA and others) and substrates. The result is reduced substrate and component gene expression in the absence of the secretory machinery, preventing substrate accumulation (20, 22). EspM also directly or indirectly regulates additional genes in *M. marinum* (22). In the presence of the ESX-1 system, WhiB6 activates the expression of the ESX-1 substrate genes and other genes, allowing production of substrates during active secretion (20, 23, 24).

The presence of EspM masked a role for EspN-dependent regulation of ESX-1 genes *in vitro*. In the absence of EspM, EspN is essential for ESX-1 activity. We showed that EspN is required for expression of the *espE* gene and contributes to the expression of the *eccA* gene, allowing optimal production of the ESX-1 membrane complex and secretion of the ESX-1 substrates. Unlike deletion of *espM* or *whiB6* (20, 22), deletion of *espN* attenuated the cytolytic activity and growth of *M. marinum* during macrophage infection, suggesting that, during infection, the transcriptional balance regulating the ESX-1 network is altered in the absence of EspN. Moreover, deletion of both *espM* and *espN* from *M. marinum* resulted in a loss of cytolytic activity during macrophage infection. EspN was likewise necessary for robust infection in a zebrafish *in vivo* model. Together, these data demonstrate that the ESX-1 transcriptional network is required during infection and identify EspN as a novel and critical regulator of this network.

Our data demonstrate that the ESX-1 system is differentially regulated during *in vitro* growth and in the host. We propose that under *in vitro* growth conditions, EspM represses expression of the ESX-1 system. Therefore, the molecular mechanisms governing protein secretion by ESX-1 have been defined primarily under conditions in which ESX-1 gene expression is repressed. Further understanding of EspN could allow us to study ESX-1 under induced conditions in the laboratory, potentially allowing the identification of new secretory components or substrates.

Our data support that EspN and EspM comprise a genetic switch that is functionally operant early in infection, including *in vivo* during zebrafish infection, where only innate immunity is present. While we do not yet understand the molecular nature of the switch, EspM and EspN likely cannot occupy the *whiB6, eccA* or *espE* promoters at the same time. Moreover, it is likely that a host-specific signal may regulate the switch including oxidative stress, pH and other phagosomal cues. Interestingly, *espN* is divergently transcribed from a putative methyltransferase gene (*MMAR_1627*) (27). We do not yet know if the genes surrounding *espN* in *M. marinum* are important for regulation of ESX-1. Interestingly, methylation has been shown to control genetic switches in bacteria (39, 40).

Expression of the ESX-1 system under *in vitro* conditions varies in diverse strains (24). Consistent with our model for a host-induced activation of ESX-1 genes by EspN, ESX-1 gene expression is upregulated in the host (41). The differential regulation of virulence in the host was suggested using transcriptomic studies in *M. marinum* in a variety of environments (42). The differential regulation of protein secretion systems under laboratory conditions and in the host is a common theme in protein secretion in bacterial pathogens. For example, some clinical *Vibrio cholerae* strains have active Type VI systems under laboratory conditions, while the pandemic strains only activate their Type VI systems in the host (43). The specialized Type III secretion systems in *Vibrio* species are regulated by bile salts (44), and by Ca^2+^ and host cell contact in *Pseudomonas* (45–47).

One interpretation of our findings is that EspN activity or *espN* expression is responsive to host-specific signals. Notably, there is evidence of infection-dependent regulation of *M. tuberculosis espN* (Rv1725c). Transcriptional profiling of bacteria from infected macrophages isolated from mouse infections revealed significant upregulation of *espN* transcript compared to broth-grown culture (48). At an early timepoint one week post infection using a mouse intravenous infection model, a TnSeq screen identified a ~2-fold decrease in representation of transposon insertions in *M. tuberculosis Rv1725c* [*P*=.0039, (49)]. However, at later timepoints (four weeks and eight weeks) this effect was no longer present (49, 50). These results and our work with *M. marinum* suggest complex regulatory circuits and genetic requirements operate at discrete stages of infection. It is also possible that EspN may be differentially localized in the mycobacterial cell in response to the host environment. SCP2 domains mediate interaction with membranes rich in acidic phospholipids (51, 52), and may serve to mediate EspN interaction with the mycobacterial cell membrane under specific conditions. SCP2 domains also mediate lipid transfer of sterols such as cholesterol and fatty acids (53, 54), which are an energy source for mycobacterial pathogens during infection (55, 56). Experiments focused on the C-terminal SCP-2 domain may reveal how EspN senses and responds to the host environment.

Alternatively, host-specific cues may remove the EspM repressor from the *whiB6* promoter. Our data suggests that EspM regulates EspN post-transcriptionally through its N-terminus, which contains an FHA domain. Proteins with FHA domains regulate other Gram-negative secretion systems post-transcriptionally (57–59). Our data also support that EspM may be processed *in vitro*. EspM cleavage might remove it from the promoter, similar to the cI repressor from λ phage (60). Alternatively, cleavage may liberate the N-terminus to regulate EspN activity, allowing the EspM C-terminus to bind DNA and repress gene expression.

Interestingly, EspN is required during infection while EspM is dispensable. EspN-dependent regulation of ESX-1 may be essential for virulence. Alternatively, EspN could regulate additional genes important for mycobacterial pathogenesis. Defining the regulatory targets of EspN, and how the EspM and EspN regulons compare, will likely require measuring global gene expression during specific stages of infection.

We were surprised that the overexpression or deletion of *espN* in the absence of *espM* resulted in the same phenotypes in *M. marinum*. It is well-established that genetic overexpression can result in loss of function phenotypes through a variety of molecular mechanisms (32). For example, *espN* overexpression could result in EspN aggregation or mislocalization. Transcription factor multimerization is well described as a regulatory mechanism in bacterial gene expression (60, 61). However, in higher organisms, transcription factors can form aggregate-like bodies that serve as functional regulatory mechanisms which mimic loss of function (62). If EspN forms multimeric aggregates when the *espN* gene is overexpressed, aggregation may prevent the interaction with DNA that mediates transcriptional activation. Alternatively, EspN may be mislocalized under overexpression conditions. Regardless of this mechanism, the same *M. marinum* strain (Δ*espN*/p*espN*) exhibited a loss of *espN* function under laboratory conditions and complementation of *espN* function in macrophages and animals. These data further support that EspN activity responds to host specific signals.

Other transcription factors regulate the ESX-1 system; however, to our knowledge, EspN is the first ESX-1 regulator that specifically functions in the host. The known transcription factors regulate ESX-1 substrates, not the ESX-1 components. Several transcription factors regulate *whiB6* and the *espACD* operon which encodes secreted substrates essential for ESX-1 secretion in *M. tuberculosis* (63–66). PhoPR and MprAB regulate *whiB6* expression directly and through the EspR regulator (67–71), in response to pH or stress, both of which are important for survival in the acidified phagosome (72–74). However, the *espACD* operon is dispensable for ESX-1 secretion and pathogenesis in *M. marinum*, and regulation of *whiB6* by PhoPR has not been reported (11, 75). Although this may suggest divergence in ESX-1 regulation between *M. marinum* and in *M. tuberculosis*, the ESX-1 transcriptional network is conserved (20, 21, 76). Widespread ESX-1-dependent gene expression has been reported in both mycobacterial species (20, 21, 76). Moreover, we showed that EspM and the regulatory substrates, EspE and EspF, are functionally conserved in *M. tuberculosis* (22, 33). Future research will be aimed at defining the ESX-1 transcriptional network in *M. tuberculosis*.

Overall, this study further defined the regulatory network underlying control of the ESX-1 secretion system and demonstrated its importance during infection. We have, for the first time, identified an infection-dependent transcriptional activator responsible for regulating both the ESX-1 components and substrates. Our study will serve as a foundation for understanding the molecular complexities of ESX-1 regulation in the host. We are now poised to define the molecular mechanisms underlying how the ESX-1 system senses and responds to a changing host environment

## Materials and Methods

Bacterial strains were derived from the *M. marinum* M parental strain (ATCC BAA-535), and were maintained as previously described (20, 22, 33, 77). Nomenclature follows the conventions proposed by Bitter et al (29). Genetic deletions were performed using allelic exchange as previously described (20, 22, 33, 77, 78). Hemolytic activity was measured against sRBCs as described previously (20, 22, 33, 77). Cell-associated and secreted mycobacterial proteins were isolated and analyzed as described in (77, 79). Protein levels were measured using western blot analysis. RNA extraction was performed from *M. marinum* using the Qiagen RNeasy kit, followed by qRT-PCR relative to the levels of *sigA* as described previously (22, 35). Macrophage (RAW 264.7) cytotoxicity was measured using Ethidium homodimer uptake following infection by *M. marinum* as described in (34, 77). Mycobacterial CFUs were performed similarly to (79). Protein modeling was performed using Robetta and Pfam as indicated. Zebrafish larvae infections with *M. marinum* were performed and bacterial burden was measured using fluorescent pixel counts as (37). Statistical analysis was performed using GraphPad Prism v. 9 or R within the latest version of R Studio IDE. Detailed methods are in the Supplementary Material.

## Acknowledgments

K.R.N is supported by an Eck Institute for Global Health Fellowship and an Arthur J. Schmitt Fellowship. P.A.C. is supported by the National Institutes of Health under award numbers AI156229, AI106872, AI149147, and AI149235. Support for D.M.T. through NIH award numbers AI130236, AI125517, and AI166304. We thank the Champion Lab for their critical reading of this manuscript. We thank Dr. Matthew Champion for assistance visualizing the data in Figure 1B. The content of this article is solely our responsibility and does not necessarily represent the official views of the National Institutes of Health.

## References

1. E. R. Green, J. Mecsas, Bacterial Secretion Systems: An Overview. Microbiol Spectr 4 (2016).

2. K. R. Nicholson, P. A. Champion, Bacterial secretion systems: Networks of pathogenic regulation and adaptation in mycobacteria and beyond. PLoS Pathog 18, e1010610 (2022).

3. P. S. Manzanillo, M. U. Shiloh, D. A. Portnoy, J. S. Cox, Mycobacterium Tuberculosis Activates the DNA-Dependent Cytosolic Surveillance Pathway within Macrophages. Cell Host Microbe 11, 469–480 (2012).

4. R. Simeone et al., Phagosomal rupture by Mycobacterium tuberculosis results in toxicity and host cell death. PLoS Pathog 8, e1002507 (2012).

5. D. Houben et al., ESX-1-mediated translocation to the cytosol controls virulence of mycobacteria. Cell Microbiol 14, 1287–1298 (2012).

6. D. M. Tobin, L. Ramakrishnan, Comparative pathogenesis of Mycobacterium marinum and Mycobacterium tuberculosis. Cell Microbiol 10, 1027–1039 (2008).

7. S. A. Stanley, S. Raghavan, W. W. Hwang, J. S. Cox, Acute infection and macrophage subversion by Mycobacterium tuberculosis require a specialized secretion system. Proc Natl Acad Sci U S A 100, 13001–13006 (2003).

8. L. Y. Gao et al., A mycobacterial virulence gene cluster extending RD1 is required for cytolysis, bacterial spreading and ESAT-6 secretion. Mol Microbiol 53, 1677–1693 (2004).

9. K. M. Guinn et al., Individual RD1-region genes are required for export of ESAT-6/CFP-10 and for virulence of Mycobacterium tuberculosis. Mol Microbiol 51, 359–370 (2004).

10. E. N. Houben et al., Composition of the type VII secretion system membrane complex. Mol Microbiol 86, 472–484 (2012).

11. R. M. Cronin, M. J. Ferrell, C. W. Cahir, M. M. Champion, P. A. Champion, Proteo-genetic analysis reveals clear hierarchy of ESX-1 secretion in Mycobacterium marinum. Proc Natl Acad Sci U S A 119, e2123100119 (2022).

12. W. H. Conrad et al., Mycobacterial ESX-1 secretion system mediates host cell lysis through bacterium contact-dependent gross membrane disruptions. Proc Natl Acad Sci U S A 114, 1371–1376 (2017).

13. F. Sayes et al., Multiplexed Quantitation of Intraphagocyte Mycobacterium tuberculosis Secreted Protein Effectors. Cell Rep 23, 1072–1084 (2018).

14. R. Wassermann et al., Mycobacterium tuberculosis Differentially Activates cGAS- and Inflammasome-Dependent Intracellular Immune Responses through ESX-1. Cell Host Microbe 17, 799–810 (2015).

15. R. O. Watson, P. S. Manzanillo, J. S. Cox, Extracellular M. tuberculosis DNA Targets Bacteria for Autophagy by Activating the Host DNA-Sensing Pathway. Cell 150, 803–815 (2012).

16. J. Smith et al., Evidence for pore formation in host cell membranes by ESX-1-secreted ESAT-6 and its role in Mycobacterium marinum escape from the vacuole. Infect Immun 76, 5478–5487 (2008).

17. N. van der Wel et al., M. tuberculosis and M. leprae translocate from the phagolysosome to the cytosol in myeloid cells. Cell 129, 1287–1298 (2007).

18. E. D. Brutinel, T. L. Yahr, Control of gene expression by type III secretory activity. Curr Opin Microbiol 11, 128–133 (2008).

19. S. N. Joslin, D. R. Hendrixson, Activation of the Campylobacter jejuni FlgSR two-component system is linked to the flagellar export apparatus. J Bacteriol 191, 2656–2667 (2009).

20. R. E. Bosserman et al., WhiB6 regulation of ESX-1 gene expression is controlled by a negative feedback loop in Mycobacterium marinum. Proc Natl Acad Sci U S A 10.1073/pnas.1710167114 (2017).

21. A. M. Abdallah et al., Integrated transcriptomic and proteomic analysis of pathogenic mycobacteria and their esx-1 mutants reveal secretion-dependent regulation of ESX-1 substrates and WhiB6 as a transcriptional regulator. PLoS One 14, e0211003 (2019).

22. K. G. Sanchez et al., EspM Is a Conserved Transcription Factor That Regulates Gene Expression in Response to the ESX-1 System. mBio 11 (2020).

23. Z. Chen et al., Mycobacterial WhiB6 Differentially Regulates ESX-1 and the Dos Regulon to Modulate Granuloma Formation and Virulence in Zebrafish. Cell Rep 16, 2512–2524 (2016).

24. L. Solans et al., A specific polymorphism in Mycobacterium tuberculosis H37Rv causes differential ESAT-6 expression and identifies WhiB6 as a novel ESX-1 component. Infect Immun 82, 3446–3456 (2014).

25. R. D. Finn et al., Pfam: the protein families database. Nucleic Acids Res 42, D222–230 (2014).

26. C. UniProt, UniProt: the universal protein knowledgebase in 2021. Nucleic Acids Res 49, D480–D489 (2021).

27. A. Kapopoulou, J. M. Lew, S. T. Cole, The MycoBrowser portal: a comprehensive and manually annotated resource for mycobacterial genomes. Tuberculosis (Edinb) 91, 8–13 (2011).

28. S. A. Shiryev, J. S. Papadopoulos, A. A. Schaffer, R. Agarwala, Improved BLAST searches using longer words for protein seeding. Bioinformatics 23, 2949–2951 (2007).

29. W. Bitter et al., Systematic genetic nomenclature for type VII secretion systems. PLoS Pathog 5, e1000507 (2009).

30. T. Parish, N. G. Stoker, Use of a flexible cassette method to generate a double unmarked Mycobacterium tuberculosis tlyA plcABC mutant by gene replacement. Microbiology 146 (Pt 8), 1969–1975 (2000).

31. V. J. van Winden et al., Mycosins Are Required for the Stabilization of the ESX-1 and ESX-5 Type VII Secretion Membrane Complexes. MBio 7 (2016).

32. G. Prelich, Gen. overexpression. uses, mechanisms, and interpretation. Genetics 190, 841–854 (2012).

33. A. E. Chirakos, K. R. Nicholson, A. Huffman, P. A. Champion, Conserved ESX-1 substrates EspE and EspF are virulence factors that regulate gene expression. Infect Immun 10.1128/IAI.00289-20 (2020).

34. E. A. Williams et al., A Nonsense Mutation in Mycobacterium marinum That Is Suppressible by a Novel Mechanism. Infect Immun 85 (2017).

35. K. G. Sanchez, R. J. Prest, K. R. Nicholson, K. V. Korotkov, P. A. Champion, Functional Analysis of EspM, an ESX-1-Associated Transcription Factor in Mycobacterium marinum. J Bacteriol 10.1128/jb.00233-22, e0023322 (2022).

36. G. M. Kennedy, G. C. Hooley, M. M. Champion, F. M. Medie, P. A. Champion, A novel ESX-1 locus reveals that surface associated ESX-1 substrates mediate virulence in Mycobacterium marinum. J Bacteriol 196, 1877–1888 (2014).

37. K. Takaki, J. M. Davis, K. Winglee, L. Ramakrishnan, Evaluation of the pathogenesis and treatment of Mycobacterium marinum infection in zebrafish. Nat Protoc 8, 1114–1124 (2013).

38. J. W. Saelens et al., An ancestral mycobacterial effector promotes dissemination of infection. Cell https://doi.org/10.1016/j.cell.2022.10.019 (2022).

39. H. N. Lim, A. van Oudenaarden, A multistep epigenetic switch enables the stable inheritance of DNA methylation states. Nat Genet 39, 269–275 (2007).

40. I. Cota et al., OxyR-dependent formation of DNA methylation patterns in OpvABOFF and OpvABON cell lineages of Salmonella enterica. Nucleic Acids Res 44, 3595–3609 (2016).

41. P. Aiewsakun et al., Transcriptional response to the host cell environment of a multidrug-resistant Mycobacterium tuberculosis clonal outbreak Beijing strain reveals its pathogenic features. Sci Rep 11, 3199 (2021).

42. E. M. Weerdenburg et al., Genome-wide transposon mutagenesis indicates that Mycobacterium marinum customizes its virulence mechanisms for survival and replication in different hosts. Infect Immun 83, 1778–1788 (2015).

43. S. T. Miyata et al., The Vibrio Cholerae Type VI Secretion System: Evaluating its Role in the Human Disease Cholera. Front Microbiol 1, 117 (2010).

44. P. Li et al., Bile salt receptor complex activates a pathogenic type III secretion system. Elife 5 (2016).

45. E. A. William McMackin, L. Djapgne, J. M. Corley, T. L. Yahr, Fitting Pieces into the Puzzle of Pseudomonas aeruginosa Type III Secretion System Gene Expression. J Bacteriol 201 (2019).

46. D. W. Frank, The exoenzyme S regulon of Pseudomonas aeruginosa. Mol Microbiol 26, 621–629 (1997).

47. J. Kim et al., Factors triggering type III secretion in Pseudomonas aeruginosa. Microbiology (Reading) 151, 3575–3587 (2005).

48. D. Pisu, L. Huang, J. K. Grenier, D. G. Russell, Dual RNA-Seq of Mtb-Infected Macrophages In Vivo Reveals Ontologically Distinct Host-Pathogen Interactions. Cell Rep 30, 335–350 e334 (2020).

49. C. M. Sassetti, E. J. Rubin, Genetic requirements for mycobacterial survival during infection. Proc Natl Acad Sci U S A 100, 12989–12994 (2003).

50. C. M. Smith et al., Host-pathogen genetic interactions underlie tuberculosis susceptibility in genetically diverse mice. Elife 11 (2022).

51. H. Huang, J. M. Ball, J. T. Billheimer, F. Schroeder, The sterol carrier protein-2 amino terminus: a membrane interaction domain. Biochemistry 38, 13231–13243 (1999).

52. E. S. Liedhegner, C. D. Vogt, D. S. Sem, C. W. Cunningham, C. J. Hillard, Sterol carrier protein-2: binding protein for endocannabinoids. Mol Neurobiol 50, 149–158 (2014).

53. H. Y. Lin, M. Y. Pang, M. G. Feng, S. H. Ying, A peroxisomal sterol carrier protein 2 (Scp2) contributes to lipid trafficking in differentiation and virulence of the insect pathogenic fungus Beauveria bassiana. Fungal Genet Biol 158, 103651 (2022).

54. M. Galano, S. Venugopal, V. Papadopoulos, Role of STAR and SCP2/SCPx in the Transport of Cholesterol and Other Lipids. Int J Mol Sci 23 (2022).

55. A. K. Pandey, C. M. Sassetti, Mycobacterial persistence requires the utilization of host cholesterol. Proc Natl Acad Sci U S A 105, 4376–4380 (2008).

56. W. Lee, B. C. VanderVen, R. J. Fahey, D. G. Russell, Intracellular Mycobacterium tuberculosis exploits host-derived fatty acids to limit metabolic stress. J Biol Chem 288, 6788–6800 (2013).

57. J. S. Lin et al., TagF-mediated repression of bacterial type VI secretion systems involves a direct interaction with the cytoplasmic protein Fha. J Biol Chem 10.1074/jbc.RA117.001618 (2018).

58. J. S. Lin et al., Fha interaction with phosphothreonine of TssL activates type VI secretion in Agrobacterium tumefaciens. PLoS Pathog 10, e1003991 (2014).

59. M. Pallen, R. Chaudhuri, A. Khan, Bacterial FHA domains: neglected players in the phospho-threonine signalling game? Trends Microbiol 10, 556–563 (2002).

60. A. D. Johnson et al., lambda Repressor and cro--components of an efficient molecular switch. Nature 294, 217–223 (1981).

61. A. Ramalinga, J. L. Danger, N. Makthal, M. Kumaraswami, P. Sumby, Multimerization of the Virulence-Enhancing Group A Streptococcus Transcription Factor RivR Is Required for Regulatory Activity. J Bacteriol 199 (2017).

62. Y. S. Huoh et al., Dual functions of Aire CARD multimerization in the transcriptional regulation of T cell tolerance. Nat Commun 11, 1625 (2020).

63. J. M. Chen et al., EspD Is Critical for the Virulence-Mediating ESX-1 Secretion System in Mycobacterium tuberculosis. J Bacteriol 194, 884–893 (2012).

64. P. A. Champion, M. M. Champion, P. Manzanillo, J. S. Cox, ESX-1 secreted virulence factors are recognized by multiple cytosolic AAA ATPases in pathogenic mycobacteria. Mol Microbiol 73, 950–962 (2009).

65. S. M. Fortune et al., Mutually dependent secretion of proteins required for mycobacterial virulence. Proc Natl Acad Sci U S A 102, 10676–10681 (2005).

66. J. A. MacGurn, S. Raghavan, S. A. Stanley, J. S. Cox, A non-RD1 gene cluster is required for Snm secretion in Mycobacterium tuberculosis. Mol Microbiol 57, 1653–1663 (2005).

67. G. Cao et al., EspR, a regulator of the ESX-1 secretion system in Mycobacterium tuberculosis, is directly regulated by the two-component systems MprAB and PhoPR. Microbiology 161, 477–489 (2015).

68. S. B. Walters et al., The Mycobacterium tuberculosis PhoPR two-component system regulates genes essential for virulence and complex lipid biosynthesis. Mol Microbiol 60, 312–330 (2006).

69. V. Anil Kumar et al., EspR-dependent ESAT-6 Protein Secretion of Mycobacterium tuberculosis Requires the Presence of Virulence Regulator PhoP. J Biol Chem 291, 19018–19030 (2016).

70. B. Blasco et al., Functional dissection of intersubunit interactions in the EspR virulence regulator of Mycobacterium tuberculosis. J Bacteriol 196, 1889–1900 (2014).

71. S. Raghavan, P. Manzanillo, K. Chan, C. Dovey, J. S. Cox, Secreted transcription factor controls Mycobacterium tuberculosis virulence. Nature 454, 717–721 (2008).

72. B. K. Johnson et al., The Carbonic Anhydrase Inhibitor Ethoxzolamide Inhibits the Mycobacterium tuberculosis PhoPR Regulon and Esx-1 Secretion and Attenuates Virulence. Antimicrob Agents Chemother 59, 4436–4445 (2015).

73. J. Gonzalo-Asensio et al., Evolutionary history of tuberculosis shaped by conserved mutations in the PhoPR virulence regulator. Proc Natl Acad Sci U S A 111, 11491–11496 (2014).

74. J. J. Baker, B. K. Johnson, R. B. Abramovitch, Slow growth of Mycobacterium tuberculosis at acidic pH is regulated by phoPR and host-associated carbon sources. Mol Microbiol 94, 56–69 (2014).

75. A. E. Chirakos, A. Balaram, W. Conrad, P. A. Champion, Modeling Tubercular ESX-1 Secretion Using Mycobacterium marinum. Microbiol Mol Biol Rev 84 (2020).

76. C. Sala et al., EspL is essential for virulence and stabilizes EspE, EspF and EspH levels in Mycobacterium tuberculosis. PLoS Pathog 14, e1007491 (2018).

77. R. E. Bosserman, K. R. Nicholson, M. M. Champion, P. A. Champion, A New ESX-1 Substrate in Mycobacterium marinum That Is Required for Hemolysis but Not Host Cell Lysis. J Bacteriol 201 (2019).

78. R. E. Bosserman, C. R. Thompson, K. R. Nicholson, P. A. Champion, Esx Paralogs Are Functionally Equivalent to ESX-1 Proteins but Are Dispensable for Virulence in Mycobacterium marinum. J Bacteriol 200, e00726-00717 (2018).

79. A. E. Chirakos, K. R. Nicholson, A. Huffman, P. A. Champion, Conserved ESX-1 Substrates EspE and EspF Are Virulence Factors That Regulate Gene Expression. Infect Immun 88 (2020).

80. M. Baek et al., Accurate prediction of protein structures and interactions using a three-track neural network. Science 373, 871–876 (2021).

