## Supplemental Figures and Tables for "The EspN transcription factor is an infection-dependent regulator of the ESX-1 system in *M. marinum*"

**Supporting Information for**
**The EspN transcription factor is an infection-dependent regulator of the ESX-1 system in *M. marinum***

Kathleen R. Nicholson, Rachel M. Cronin, Aruna R. Menon, Madeleine K. Jennisch, David M. Tobin, Patricia A. Champion

Patricia A. Champion


**This PDF file includes:**

Supporting Methods
Figures S1 to S6
Tables S1 to S3
SI References

### Supporting Methods

**Growth and Generation of Bacterial Strains** Mycobacterial strains were derived from the *M. marinum* M parental strain (“WT”, ATCC BAA-535). Parental strains including a 3X-FLAG epitope at the C-terminus of WhiB6 are denoted by “W6Fl” and were used for WhiB6 protein detection (1). All bacterial strains were maintained as described previously (1-4) . All liquid cultures of *M. marinum* strains were grown in Middlebrook 7H9 (Sigma Aldrich, St. Louis, MO) broth with 0.5% glycerol and 0.1% Tween-80 unless otherwise noted at 30°C. For growth on agar plates, *M. marinum* strains were struck for isolation on Middlebrook 7H11 (Sigma Aldrich, St. Louis, MO) agar supplemented with 0.5% glycerol and 0.5% glucose. Broth or agar were supplemented with 20μg/ml kanamycin (IBI Scientific, Dubuque, IA) or 50μg/ml hygromycin (Sigma-Aldrich) as needed. Strains containing integrating plasmids were grown in the absence of antibiotics in liquid media. All assays used the following estimation: 1 $OD_{600}=7.7x10^7$ cells/ml for *M. marinum*. *E. coli* strains were grown in LB (Luria-Bertani) media (VWR) with 50 μg/ml kanamycin, 200 μg/ml hygromycin, 200 μg/ml ampicillin, or 12 μg/ml tetracycline (Thermo Fisher, Waltham, MA) when needed. Cloning and plasmid propagation was performed using DH5α *E. coli* (New England Biolabs, Ipswich, MA). *E. coli* strains were grown at 37°C.

**Nomenclature**. Nomenclature in this work follows the convention set by Bitter et al. (6). ESX-1 conserved components are denoted Ecc. ESX-1 associated proteins are denoted Esp. Genes labeled with a subscript 1 are associated with the ESX-1 system.

**Generation of mycobacterial strains**. Oligonucleotide primers were purchased from Integrated DNA Technologies (IDT, Coralville, IA). Plasmids were generated using FastCloning using either *M. marinum* M or *M. tuberculosis* Erdman genomic DNA. Plasmids were introduced into *M. marinum* using electroporation as described previously (1-4, 7, 8). Strains, primers, and plasmids are listed in Supplementary Tables S1, S2, and S3. All plasmids and genetic deletions were confirmed by targeted DNA sequencing performed by the Notre Dame Genomics and Bioinformatics Facility.

**Allelic Exchange.** Allelic exchange was performed as previously published to generate mutant *M. marinum* strains (1-4, 8). Briefly, approximately 1,500 base pairs upstream and downstream of the annotated open reading frame (9) was amplified using PCR with the Phusion polymerase. The PCR amplified upstream and downstream regions were introduced into the PCR amplified p2NIL vector [Addgene plasmid number 20188; a gift from Tanya Parish (10) by three-part FastCloning (11)]. Following transformation into DH5α *E. coli* (NEB), colonies were picked from LB plates containing kanamycin (50μg/ml). Plasmids were confirmed using restriction enzyme digestion with enzymes chosen depending on the insert composition. Next, confirmed p2NIL plasmids were digested with PacI (NEB), treated with Antarctic Phosphatase (NEB), and ligated with the pGOAL19 vector (Addgene plasmid number 20190; a gift from Tanya Parish (10), as previously described (1-4). Following confirmation by restriction digestion with PacI and AflII (NEB) and targeted DNA sequencing at the Notre Dame Genomics and Bioinformatics Facility, plasmids were quantified using a Nanodrop (Thermo

Fisher). 2 µg of the confirmed plasmid was irradiated with 0.1 J/cm2 of UV light using a CL-1000 UV crosslinker (UVP). Electrocompetent *M. marinum* cells were transformed using electroporation. 500 µl of *M. marinum* competent cells were electroporated in a Gene Pulser XCell (Bio-Rad) in the presence of 2µg of the UV irradiated plasmid. Transformed *M. marinum* cells were allowed to recover overnight in 2ml of 7H9 media with 0.1% Tween-80. Following overnight recovery, cells were pelleted by centrifugation, resuspended in 200 µl of medium, and plated on 7H11 agar (Sigma) supplemented with oleic acid-albumin-dextrose-catalase (OADC), 50 µg/ml hygromycin (Corning), and 60 µg/ml 5-bromo-4-chloro-3-indolyl-β-D-galactopyranoside (X-gal) with 2% sucrose (Macron).

**Hemolysis Assays.** Sheep red blood cell (sRBC) assays were performed as previously described (1-4). Briefly, *M. marinum* strains were grown in 7H9 medium with 0.1% Tween-80. Bacteria were grown to an $OD_{600}$ between $2.0x10^8$ cells and $6.5x10^8$ cells. $5.0x10^8$ bacteria were washed three times with 1X phosphate buffered saline (PBS) and incubated with sRBCs (Hardy Diagnostics, Santa Maria, CA) at 30°C for 2h. $OD_{405}$ readings were obtained in technical triplicate using a SpectraMax plate reader.

**Protein preparation and analysis: ESX-1 secretion assays.** ESX-1 protein secretion assays were performed exactly as previously described (2, 12). *M. marinum* was cultured in 5 ml of Middlebrook 7H9 defined broth, (Sigma-Aldrich, St. Louis, MO) + 0.1% Tween-80 (Fisher Scientific, Pittsburgh, PA) for three days, then moved to 25 ml Middlebrook 7H9 + 0.1% Tween-80 for two days. *M. marinum* strains were diluted to an $OD_{600}$ of 0.8 in Sauton's broth + 0.01%

Tween-80. Following 48 hours of growth at 30°C, *M. marinum* cells were collected by centrifugation. The cells were lysed using a Biospec Mini-BeadBeater-24, and the lysate was clarified by centrifugation. The resulting proteins are the cell-associated protein fraction. Phenylmethylsulfonyl fluoride (PMSF, Roche) was added to the supernatants to a 0.1% final concentration. The supernatant was then filtered using 0.2 µm Nalgene Stericups with polyethersulfone (PES) filters to remove extraneous bacteria. Supernatants were concentrated by ultrafiltration using a 3,000-molecular-weight-cutoff (MWCO) Amicon filter (Millipore, Sartorius), yielding the secreted protein fraction. Pellets were resuspended in 1X PBS + 0.1% PMSF (Roche) and bead beaten using a Biospec Mini-BeadBeater-24 and silica beads. Pellets were clarified by centrifugation and transferred to new tubes. Mycobacterial cell-associated protein fractions were quantified using a Micro BCA Protein Assay Kit (ThermoFisher).

**Western Blotting.** All SDS-PAGE gels were loaded with 10µg of protein unless otherwise indicated. For mycobacterial protein samples, all SDS-PAGE gels were 4-20% gradients (BioRad). All antibodies were diluted in 5% nonfat dry milk in 1X PBS + 0.1% Tween-20. Rpoβ (anti-RNA polymerase beta mouse monoclonal antibody [clone: 8RB13]; VWR) was diluted 1:20,000. The following reagents were obtained through BEI resources, NIAID, NIH: polyclonal anti-*Mycobacterium tuberculosis* CFP-10 (gene Rv3874; antiserum, rabbit; NR-13801) and polyclonal anti-*Mycobacterium tuberculosis* Mpt-32 (gene Rv1860; antiserum, rabbit; NR-13807). CFP-10 was used at a 1:5,000 dilution. Mpt-32 was used at a 1:30,000 dilution. The EspE antibody (1:5,000 dilution) was obtained from Frederic

Carlsson (13). $EccCb_1$ was detected using a custom rabbit polyclonal antibody against the CDKQEFPSSEFKVKR peptide (Genscript). The $EccCb_1$ antibody was used at a 1:5,000 dilution. The C-terminally tagged WhiB6 protein was detected using a monoclonal α-FLAG M2 antibody (Millipore) at a 1:5,000 dilution. The C-terminally tagged EspM protein was detected using a mouse monoclonal α-V5 antibody (Sigma) at a 1:5,000 dilution. Horse radish peroxidase (HRP)-conjugated goat α-mouse immunoglobulin secondary antibody (Bio-Rad) was used at a 1:5000 dilution to detect the α-EsxA, α-RNAP, α-FLAG, and α-V5 antibodies. Horse radish peroxidase (HRP)-conjugated goat α-rabbit immunoglobulin secondary antibody (Bio-Rad) was used at a 1:20,000 dilution to detect the α-Mpt32 and α-EsxB antibodies. Horse radish peroxidase (HRP)-conjugated goat α-rabbit immunoglobulin secondary antibody (Bio-Rad) was used at a 1:5,000 dilution to detect α-EspE. All proteins were detected using the LumiGLO chemiluminescent substrate kit (SeraCare, Milford, MA) and X-ray film (RPI, Mt. Prospect, IL).

**RNA Extraction.** *M. marinum* was cultured in 5 ml of Middlebrook 7H9 media (Sigma Aldrich) + 0.1% Tween-80 (Fisher Scientific, Pittsburgh, PA) for three days, then moved to 25 ml Middlebrook 7H9 + 0.1% Tween-80 for two days. *M. marinum* strains were diluted to $OD_{600}$=0.8 in Sauton's broth + 0.01% Tween-80 and grown for 48h at 30°C. Bacterial culture pellets were collected by centrifuging 15 ml of Sauton's culture. These pellets were frozen. Thawed bacterial pellets were resuspended Qiagen RLT buffer (Qiagen, Hilden, Germany) supplemented with 1% β-mercaptoethanol. Lysates were generated by bead beating pellet resuspension 3 times for 30 seconds using silica beads and a Biospec Mini-BeadBeater-16 (BioSpec Products Inc., Batesville, OH, USA). Total RNA was extracted from clarified lysates using the RNeasy Mini Kit (Qiagen), according to manufacturer's instructions.

**qRT-PCR.** 500 ng-1 µg of RNA was treated with Promega RQ1 DNase (Promega) according to manufacturer instructions and supplemented with 5mM $MgCl_2$ and 10mM $CaCl_2$. 1µl of DNase treated RNA was converted to cDNA using random hexamers (IDT) and Superscript II (SSII) Reverse Transcriptase (Invitrogen) according to manufacturer's instructions. cDNA was quantified using a NanoDrop 2000 (Thermo Fisher).

qRT-PCR reactions were prepared using 250ng of cDNA mixed with SYBR Select Master Mix (Applied Biosystems, Carlsbad, CA) and 1µM of each oligonucleotide (unless noted otherwise). *sigA* was used as a housekeeping reference gene control and was detected using oligonucleotide primers sigA-F/sigA-R. All other oligonucleotide primers are listed in Supplementary Table S3. All qRT-PCR reactions were run using Applied Biosystems MicroAmp Fast 96 well plates (0.1mL) (Life Technologies). All plates were run on a QuantStudio 3 Real-Time PCR System (Thermo Fisher). Cycle conditions were as follows: 50°C for 2 min., 95°C for 10 min; 40 cycles at 95°C for 15 sec and 60°C for 1 min; a dissociation step of 95°C for 15 sec., 60°C for 1 min., 95°C for 15 sec., and 60°C for 15 sec.

All qRT-PCR reactions were analyzed using ΔΔ Ct comparisons. All qRT-PCR results were normalized to WT transcript abundance using the following equations:

$$\Delta Ct = Ct(\text{gene of interest}) - Ct(\text{housekeeping gene})$$

Then:

$$\Delta\Delta Ct = \Delta Ct(\text{treated sample}) - \Delta Ct(\text{untreated sample})$$

Then:

$$2^{-\Delta\Delta Ct} = \text{fold change}$$

**Macrophage Cytotoxicity.** RAW264.7 murine macrophages (ATCC TIB-71) were grown in high glucose, high pyruvate Dulbecco's Modified Eagle's Medium (DMEM) (Gibco, Dublin, Ireland) with 10% heat-inactivated fetal bovine serum (FBS) (Avantor, Radnor, PA). All RAW264.7 cells were cultured at 37°C under 5% $CO_2$. Macrophages were washed with sterile 1X phosphate buffered saline (PBS) pH 7.4 (Gibco, Dublin, Ireland) and passaged using cell scraping as needed.

Macrophage infections were performed as described previously (2, 14). Briefly, macrophage monolayers were seeded in 1ml DMEM + 10% FBS at $3x10^5$ in 24-well plates (Greiner Bio-One, Germany). 24h after seeding, RAW264.7 cells were infected in technical triplicate with mycobacterial strains at an MOI=4, where 1 $OD_{600}$ = $7.7x10^7$ cells. At 2 hpi, each well was treated with 100 µg/ml Gentamicin (RPI Corporation, Mt. Prospect, IL). Macrophage monolayers were washed 3X with sterile 1X PBS 6 hpi and fresh DMEM + 10% FBS was added. To assay cytotoxicity, culture media was removed 24 hpi and 250μl of EthD-1 (1μl/ml) + Calcein-AM (0.25ul/ml) (Live/Dead Viability/Cytotoxicity Kit; Life Technologies, Carlsbad, CA) solution in 1X PBS was added. Cells were incubated for 30 min at 37°C + 5% $CO_2$ and imaged using a Zeiss AxioObserver A1 inverted microscope with phase-contrast, rhodamine (red), and green fluorescent protein (GFP) filters. Ten images were obtained per well, and dead cells were quantified using ImageJ as described previously (14).

**CFU Assays.** RAW 264.7 macrophages were seeded in 1ml DMEM plus 10% FBS per well at $5x10^5$ cells/ml in a 24-well plate (Greiner Bio-One, Germany) and allowed to grow for 24h at 37°C + 5% $CO_2$. Bacteria were added at an MOI of 0.2 ($1x10^5$ cells/ml) in technical triplicate and mixed. Infections proceeded for 2h at 37°C + 5% $CO_2$. 2-hour post-infection (2hpi) entry assays were harvested. For harvesting, medium was removed and 0.5ml of sterile lysis buffer (H2O plus 0.1% [vol/vol] Tween 80) was added. Plates were then incubated at 37°C + 5% $CO_2$ for 10min before scraping the wells and pipetting up and down. Cells were diluted at 1:1000 using sterilized dilution buffer (1x PBS plus 0.05% [vol/vol] Tween 80). 50μl of the dilutions were plated on Middlebrook 7H11 plates supplemented with 10% (vol/vol) oleic acid-albumin-dextrose-catalase (OADC) and 0.5% (vol/vol) glycerol. For uninfected controls, 50μl of undiluted cells were plated. Each technical triplicate was plated in duplicate (6 plates/time point). For 24, 48, 72, and 96hpi time points, gentamycin (RPI Corporation, Mt. Prospect, IL) was added at 100μg/ml 2hpi. At 4hpi, cells were washed three times with sterile 1x PBS and 1ml fresh medium was added to each well. All time points were harvested exactly as described above. At 48hpi, wells for 72 and 96hpi time points were washed three times with sterile 1x PBS and 1ml fresh medium was added to each well. Colonies were counted following approximately 1 week-incubation at 32°C.

**Protein Modeling and Alignments.** Protein modeling was performed using Robetta, Pfam, and Alphafold as indicated (15-17). Protein sequences were obtained from Mycobrowser or Biocyc, where appropriate (9, 18).

**Zebrafish infections.** Zebrafish larvae were anaesthetized and infected at 2 days post-fertilization with 150-200 colony forming units (CFU) of each strain and assessed at 1-day post-infection and 5 days post-infection for bacterial burden using fluorescent pixel counts (19), a validated longitudinal readout of bacterial burden. 1-day post infection readouts controls for starting levels of bacterial burden. Fluorescence is enumerated over the area of the infected fish by calculating the number of pixels above background using a constant threshold. These results were analyzed using R 4.2.2 within the latest version of RStudio IDE using in-house workflows. Fold change was the ratio of fluorescence of *ΔespN* compared to WT. Graphing was similarly performed in R using ggplot2. All zebrafish husbandry and experimental procedures were performed in accordance and compliance with policies approved by the Duke University Institutional Animal Care and Use Committee (protocol A091-20-04).

**Statistical Analysis**. All statistical analyses were performed in GraphPad Prism version 9 or R 4.2.2 within the latest version of RStudio IDE. Statistical significance was determined by performing an ordinary one-way or two-way ANOVA followed by either a Dunnett's, Tukey's, or Sidak's multiple comparisons test. P-values for individual experiments are noted in the figure legend and in the text. All experiments were conducted on at least three biological replicates, each with technical triplicates where available.

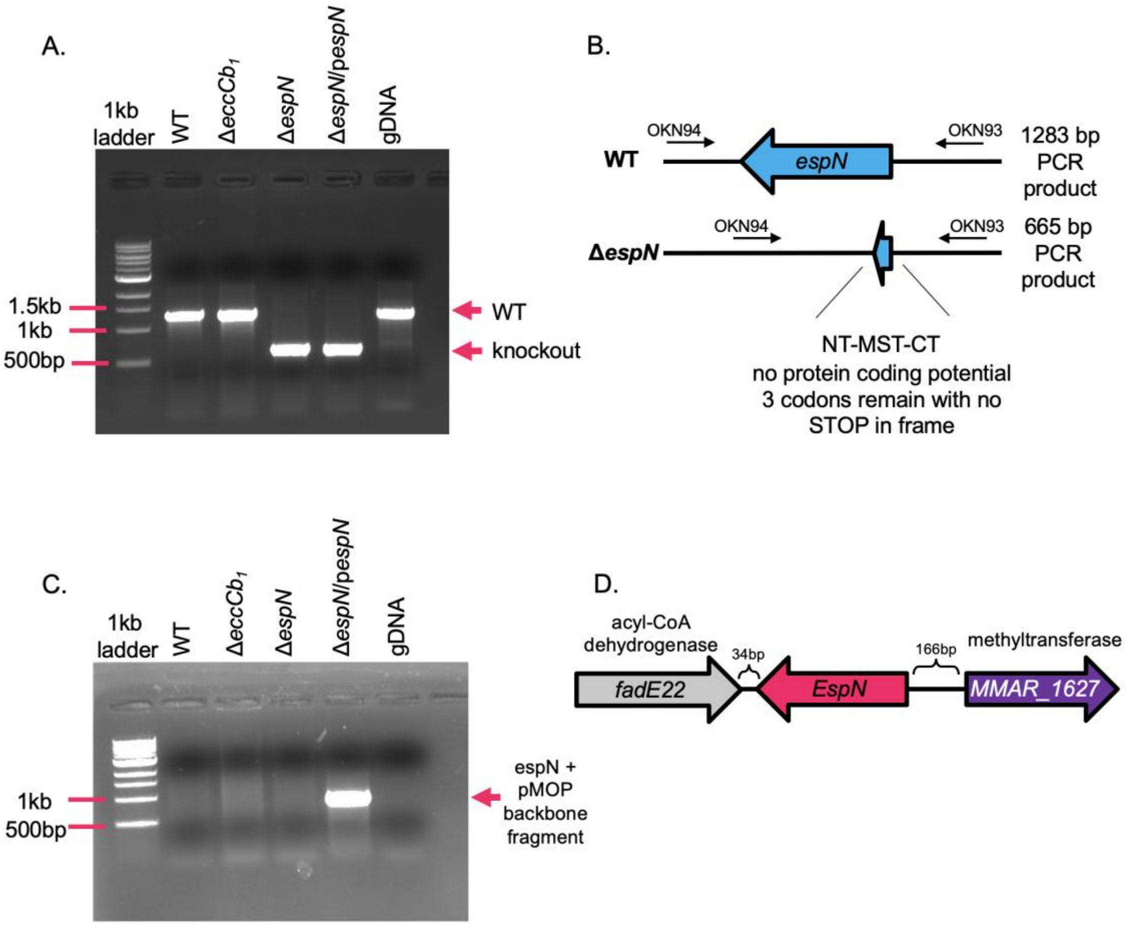


**Fig. S1. Genotypic confirmation of *espN* deletion and complementation. A.** PCR of upstream and downstream of the *espN* gene using OKN93 and OKN94 primers to confirm genetic deletion in *M. marinum* strains. WT = 1283 bp; Δ*espN* = 665 bp. *M. marinum* genomic DNA (gDNA) serves as a negative control. **B.** Schematic of the *espN* deletion using allelic exchange. **C.** PCR of p*espN* plasmid in *M. marinum* strains using OKN139 and MOPF primers. A 775 bp band indicates the p*espN* is present. *M. marinum* gDNA serves as a negative control. **D.** Schematic of the genetic locus including *espN* in *M. marinum.*

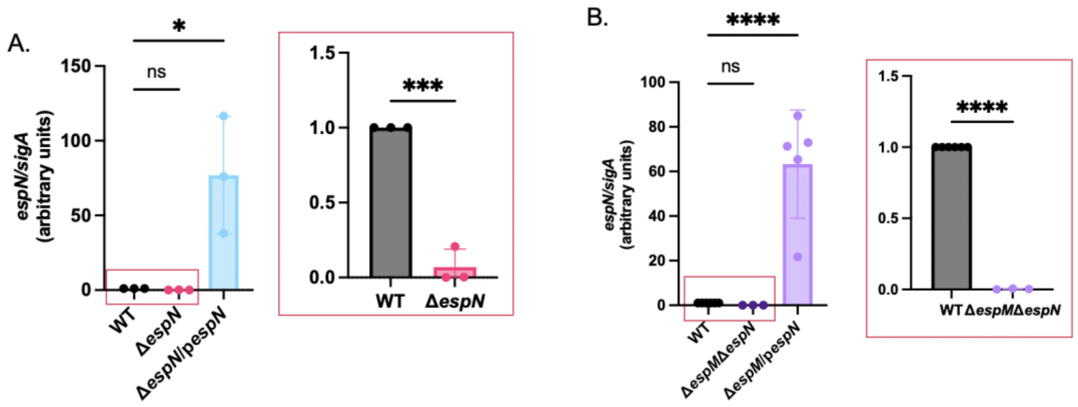


**Fig. S2. EspN expression in deletion and overexpression *M. marinum* strains.** A. Relative qRT analysis of *espN* in *M. marinum* strains compared to *sigA* transcript levels. Statistical analysis was performed using a one way-ordinary ANOVA (*P*=.0092), followed by a Dunnett's multiple comparison test. * *P*=.0114. Inset: Zoom in of the comparison between the WT and Δ*espN* strain. The levels of *espN* were compared using an unpaired student's t-test. *** *P*=.0002. B. Relative qRT analysis of *espN* in *M. marinum* strains compared to *sigA* transcript levels. Statistical analysis was performed using a one way-ordinary ANOVA (*P*<.0001), followed by a Dunnett's multiple comparison's test. **** *P*<0.0001. Inset: Comparison between the WT and Δ*espM*Δ*espN* strains. The levels of *espN* were compared using an unpaired student's t-test. **** *P*<0.0001.

A

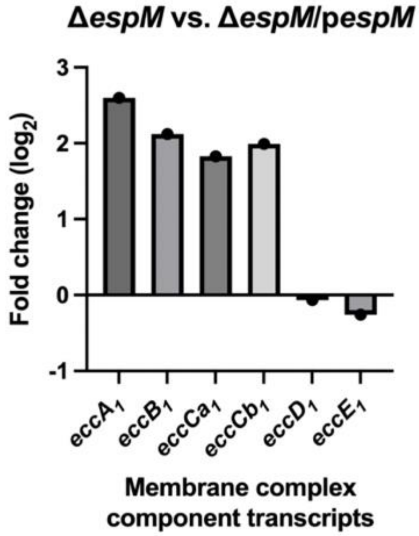


B

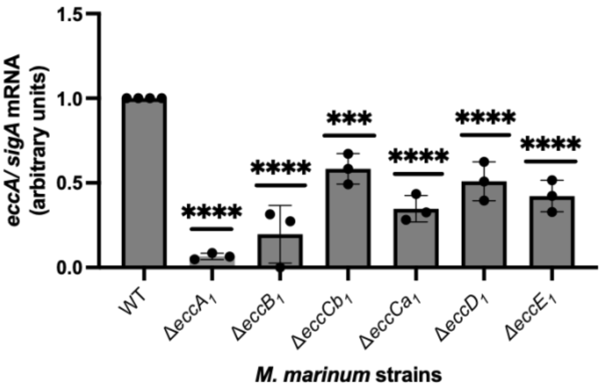


**Fig. S3. ESX-1 membrane complex transcripts are regulated by ESX-1. A.** $Log_2$ fold-change of ESX-1 membrane complex component transcripts from RNA sequencing data published in Sanchez et al (3). The *eccA-eccCb*$_1$ transcripts were significantly upregulated in the Δ*espM* strain compared to the overexpression strain (*P = 1.07e-23, 1.11e-19, 3.99e-15, 2.13e-16*). The *eccD*$_1$ and *eccE*$_1$ transcripts were not significantly different between the two strains by RNA sequencing. **B.** Relative qRT analysis of *eccA* transcript in *M. marinum* strains compared to *sigA* transcript levels. Statistical analysis was performed using a one way-ordinary ANOVA *(P*=.0001), followed by a Tukey's multiple comparison test. *** *P*=.0005, **** *P*<.0001.

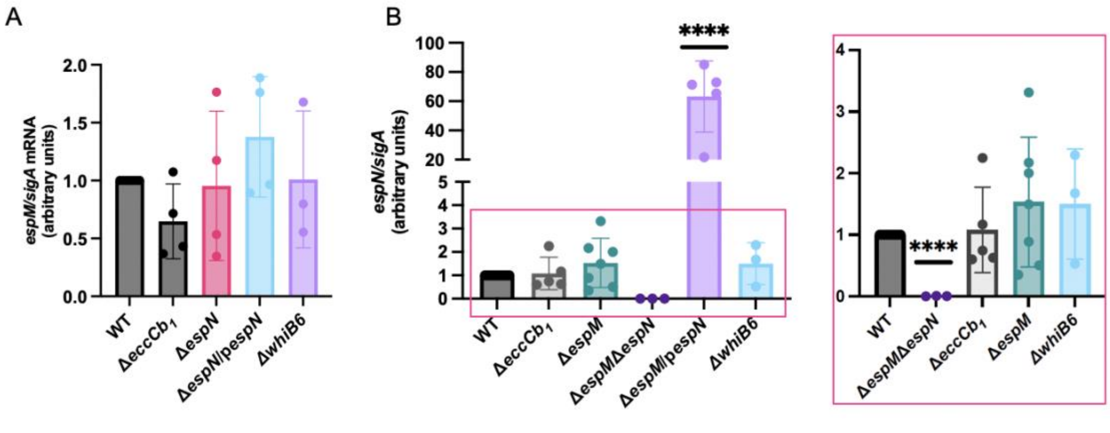


**Fig. S4: EspM and EspN do not regulate each other transcriptionally in *M. marinum* under laboratory conditions**. qRT PCR on total RNA extracted from *M. marinum* following growth *in vitro* measuring **A.** *espM* transcript or **B.** *espN* transcript relative to *sigA*. None of the measured changes of the *espM* transcript in panel A are significantly different from the WT strain based on a one-way ordinary ANOVA. In panel B, significance was determined using a one-way ordinary ANOVA (*P<.0001*), followed by a Dunnett's multiple comparison test. **** *P*<.0001 compared to the WT strain. In the inset, the Δ*espM*/*pespN* strain was excluded. In the inset, an unpaired t-test was performed between the WT and the Δ*espM*Δ*espN* and WT strains (*P*<.0001).

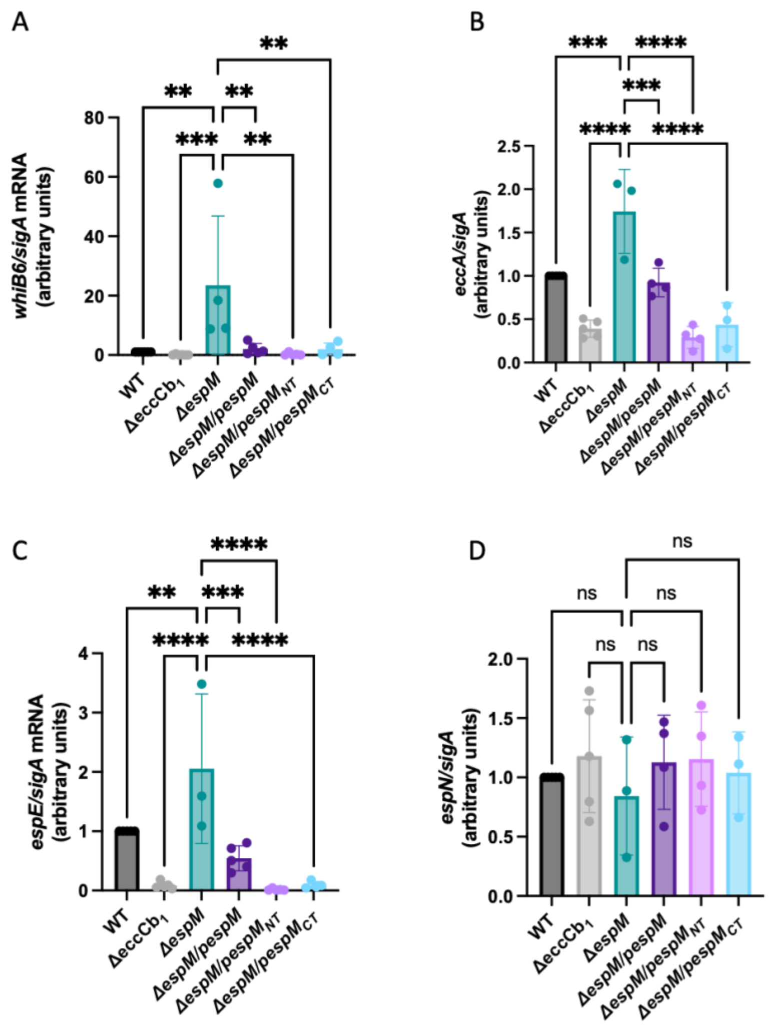


**Fig. S5: Expression of EspM$_{NT}$ impacts ESX-1 gene expression.** qRT-PCR analysis of the A) *whiB6,* B) *eccA,* C) *espE* and D)*espN* relative to *sigA.* Each data point is the average of three technical replicates. Statistical analysis was performed using a one-way ordinary ANOVA followed by a Dunnett's post-hoc test vs the Δ*espM* strain. A) ANOVA *P*<.0001, ** (vs WT, *P*=.0014, vs Δ*espM/pespM, P*=.0032, vs Δ*espM/pespM*$_{NT}$*, P*=.0015, Δ*espM/pespM*$_{CT}$*, P*=.0050), *** *P* =.0009. B) ANOVA *P*<.0001, *** (vs WT, *P*=.0005, vs Δ*espM/pespM, P*=.0003), **** *P* <.0001, C) ANOVA *P*<.0001, ** *P*=.0068, *** *P*=.0002, **** *P* <.0001, D) ANOVA *P* value was not significant. These data are represented as the heat map in Figure 4B.

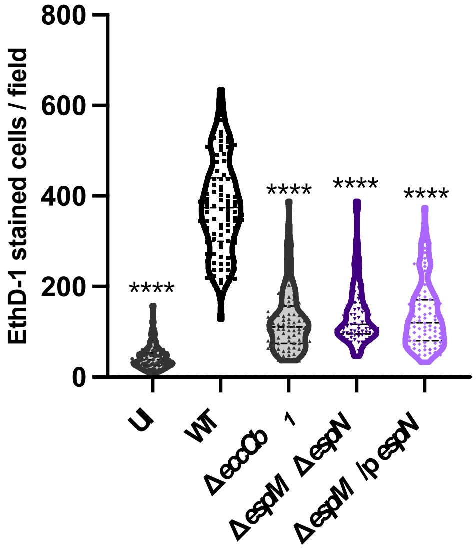


**Fig. S6: The loss of EspM and EspN attenuate *M. marinum* in a macrophage model of infection.** Macrophage cytolysis (RAW 264.7 cells) as measured by EthD-1 staining 24h post infection with *M. marinum* strains at an MOI of 4. Statistical analysis was performed using a one-way ANOVA followed by a Dunnett's multiple comparisons test relative to the WT strain. (**** $P$<.0001) Each dot represents the number of EthD-1-stained cells in a single field. A total of 10 fields were counted using ImageJ for each well. Processing of 3 wells was performed for each biological replicate. A total of 90 fields were counted for each strain. UI: uninfected.

**Table S1. Strains used in this study.**

| Name | Genotype | Reference |
|---|---|---|
| *M. marinum* M strain | Wild type; parent for all strains | ATCC BAA-535 |
| Δ*whiB6* | M with deletion of the *whiB6* (*MMAR_5437) gene* | (1) |
| *whiB6*-FL | M with *whiB6* allele tagged with 3X-FLAG at C-terminus, at *whiB6* locus | (1) |
| *whiB6*-FLΔ$eccCb_1$ | *whiB6*-FL with deletion of the M with deletion of the $eccCb_1$ *(MMAR_5446)* gene | (1) |
| *whiB6*-FLΔ*espN* | *whiB6*-FL with deletion of the M with deletion of the *espN (MMAR_1626)* gene | This study |
| *whiB6*-FLΔ*espN*/p*espN* | *whiB6*-FLΔ*espN* with an integrating plasmid expressing *espN* from the Mycobacterial Optimal Promoter | This study |
| *whiB6*-FLΔ*espM* | *whiB6*-FL with deletion of the M with deletion of the *espM (MMAR_5438)* gene | (3) |
| *whiB6*-FLΔ*espM*Δ*espN* | *whiB6*-FLΔ*espM* with deletion of the *espN (MMAR_1626)* gene | This study |
| *whiB6*-FLΔ*espM*/p*espN* | *whiB6*-FLΔ*espM* with an integrating plasmid expressing *espN* from the Mycobacterial Optimal Promoter | This study |
| *whiB6*-FLΔ*espM*/p*espM-V5* | *whiB6*-FLΔ*espM* with an integrating plasmid expressing *espM* with a C-terminal V5 tag from the Mycobacterial Optimal Promoter | (20) |
| *whiB6*-FLΔ*espM*/p$espM_{CT}$-*V5* | *whiB6*-FLΔ*espM* with an integrating plasmid expressing the C-terminal half (AA 127-363) of *espM* with a C-terminal V5 tag from the Mycobacterial Optimal Promoter | (20) |
| *whiB6*-FLΔ*espM*/p$espM_{NT}$-*V5* | *whiB6*-FLΔ*espM* with an integrating plasmid expressing the N-terminal half (AA 1-133) of *espM* with a C-terminal V5 tag from the Mycobacterial Optimal Promoter | This study |

**Table S2. List of plasmids used in this study**

| Name | Genotype | Reference |
|---|---|---|
| p2NIL | *kan*$^R$, *amp*$^R$, *oriE*; Parental vector for allelic exchange | (10) |
| pGOAL19 | *amp*$^R$, GOAL cassette includes *hygR*, *lacZ*, *sacB*, *oriE*; Parental vector for allelic exchange | (10) |
| p2NILΔ*espN*GOAL | *M. marinum espN* flanking regions. *kan*$^R$, *hyg*$^R$, *lacZ, sacB* | This study |
| p*espN* | *espN* from *M. marinum* behind the pMOP promoter, *hyg*$^R$. *attB* integration | This study |
| pMSP12*mCerulean* | *mCerulean* gene expressed behind the pMSP12 promoter. *kan*$^R$. Episomal. | (21) |
| p*espM*-V5 | *espM (MMAR_5438)* from *M. marinum* behind the pMOP promoter, *hyg*$^R$. *attB* integration. V5 C-terminal tag. | (20) |
| p*espM*$_{NT}$-V5 | Amino acids 1-133 from *espM (MMAR_5438)* from *M. marinum* behind the pMOP promoter, *hyg*$^R$. *attB* integration. V5 C-terminal tag. | (20) |
| p*espM*$_{CT}$-V5 | Amino acids 127-363 from *espM (MMAR_5438)* from *M. marinum* behind the pMOP promoter, *hyg*$^R$. *attB* integration. V5 C-terminal tag. | This study |

**Table S3: List of oligonucleotide primers used in this study.**

| Name | Sequence 5' → 3' | Application and Reference |
|---|---|---|
| OKN88 | cgtggtgtcacgctcgtGACATCACCGG GCTCAACAAC | Primer pair (A&B, upstream arm) to delete *MMAR_1626* using p2NIL. This study. |
| OKN89 | CGGGCCTTAAGATGTGGACATC GCGCAAGACCC | |
| OKN90 | CCACATCTTAAGGCCCGCGAGG CGGATGAC | Primer pair (C&D, downstream arm) to delete *MMAR_1626* using p2NIL. This study. |
| OKN91 | gcagtcaggcaccgtATCGGATTCACC TGGGAGCATGAC | |
| OKN93 | TATCGCCACGTTGCCAGATCC | Δ*MMAR_1626* F and R genotyping primers This study. |
| OKN94 | TCAAGCTGGCCGAACACATGG | |
| OKN138 | aggagtccagccatTGCGCGATGTCC ACATACAGC | F and R primers to generate *MMAR_1626* insert for FastCloning into pMOP. This study. |
| OKN139 | gcctgagcggtcccgactagtATCCGCCT CGCGGGCTCAG | |
| sigA-F | TCGAGGTGATCAACAAGCTG | *sigA* qRT primers (14) |
| sigA-R | TGGATCTCCAGCACCTTCTC | |
| olc 208 | GACGGCGTCTACAAGGTCTG | *espE* qRT primers (4) |
| olc 209 | CCGGAATGTTCGGGAGTAGG | |
| ORS225 | AGATTCCGCTGGGCGTTTGC | *whiB6* qRT primers (1) |
| ORS226 | TCTGCCAGCGACCGAAGTTG | |
| 5438FqRT | CGTCACCAACAGCCCAAACG | *espM* qRT primers (3) |
| 5438RqRT | CTGCGCTGACTGATGTCGAG | |
| ORS152 | GCCTTCGTCAGTGAGTTTCC | *eccCb$_1$* qRT primers (1) |
| ORS153 | TGGGCTGCGATTTGAGCTAC | |
| OKNq24 | TTGAAGGATCCGTCCTACCG | *eccA* qRT primers This study. |
| OKNq25 | CGAGTTCTTCTTGGGCTTCG | |
| OKNq46 | CCACATACAGCCAGTTCTGC | *espN* (*MMAR_1626*) qRT primers This study. |
| OKNq47 | GCCTTGGTCAACGACTTGAG | |

### SI References


1. R. E. Bosserman *et al.*, WhiB6 regulation of ESX-1 gene expression is controlled by a negative feedback loop in Mycobacterium marinum. *Proc Natl Acad Sci U S A* 10.1073/pnas.1710167114 (2017).
2. R. E. Bosserman, K. R. Nicholson, M. M. Champion, P. A. Champion, A New ESX-1 Substrate in Mycobacterium marinum That Is Required for Hemolysis but Not Host Cell Lysis. *J Bacteriol* **201** (2019).
3. K. G. Sanchez *et al.*, EspM Is a Conserved Transcription Factor That Regulates Gene Expression in Response to the ESX-1 System. *mBio* **11** (2020).
4. A. E. Chirakos, K. R. Nicholson, A. Huffman, P. A. Champion, Conserved ESX-1 substrates EspE and EspF are virulence factors that regulate gene expression. *Infect Immun* 10.1128/IAI.00289-20 (2020).
5. D. A. Daines, R. P. Silver, Evidence for multimerization of neu proteins involved in polysialic acid synthesis in Escherichia coli K1 using improved LexA-based vectors. *J Bacteriol* **182**, 5267-5270 (2000).
6. W. Bitter *et al.*, Systematic genetic nomenclature for type VII secretion systems. *PLoS Pathog* **5**, e1000507 (2009).
7. T. Parish, Electroporation of Mycobacteria. *Methods Mol Biol* **2314**, 273-284 (2021).
8. R. E. Bosserman, C. R. Thompson, K. R. Nicholson, P. A. Champion, Esx Paralogs Are Functionally Equivalent to ESX-1 Proteins but Are Dispensable for Virulence in Mycobacterium marinum. *J Bacteriol* **200**, e00726-00717 (2018).
9. A. Kapopoulou, J. M. Lew, S. T. Cole, The MycoBrowser portal: a comprehensive and manually annotated resource for mycobacterial genomes. *Tuberculosis (Edinb)* **91**, 8-13 (2011).
10. T. Parish, N. G. Stoker, Use of a flexible cassette method to generate a double unmarked Mycobacterium tuberculosis tlyA plcABC mutant by gene replacement. *Microbiology* **146 ( Pt 8)**, 1969-1975 (2000).
11. C. Li *et al.*, FastCloning: a highly simplified, purification-free, sequence- and ligation-independent PCR cloning method. *BMC Biotechnol* **11**, 92 (2011).
12. A. E. Chirakos, K. R. Nicholson, A. Huffman, P. A. Champion, Conserved ESX-1 Substrates EspE and EspF Are Virulence Factors That Regulate Gene Expression. *Infect Immun* **88** (2020).
13. F. Carlsson, S. A. Joshi, L. Rangell, E. J. Brown, Polar localization of virulence-related Esx-1 secretion in mycobacteria. *PLoS Pathog* **5**, e1000285 (2009).
14. E. A. Williams *et al.*, A Nonsense Mutation in Mycobacterium marinum That Is Suppressible by a Novel Mechanism. *Infect Immun* **85** (2017).
15. M. Baek *et al.*, Accurate prediction of protein structures and interactions using a three-track neural network. *Science* **373**, 871-876 (2021).

16. J. Jumper *et al.*, Highly accurate protein structure prediction with AlphaFold. *Nature* **596**, 583-589 (2021).
17. R. D. Finn *et al.*, Pfam: the protein families database. *Nucleic Acids Res* **42**, D222-230 (2014).
18. P. D. Karp *et al.*, The BioCyc collection of microbial genomes and metabolic pathways. *Brief Bioinform* **20**, 1085-1093 (2019).
19. K. Takaki, C. L. Cosma, M. A. Troll, L. Ramakrishnan, An in vivo platform for rapid high-throughput antitubercular drug discovery. *Cell Rep* **2**, 175-184 (2012).
20. K. G. Sanchez, R. J. Prest, K. R. Nicholson, K. V. Korotkov, P. A. Champion, Functional Analysis of EspM, an ESX-1-Associated Transcription Factor in Mycobacterium marinum. *J Bacteriol* 10.1128/jb.00233-22, e0023322 (2022).
21. J. W. Saelens *et al.*, An ancestral mycobacterial effector promotes dissemination of infection. *Cell* https://doi.org/10.1016/j.cell.2022.10.019 (2022).
